# Maintenance cost of photosynthesis sets key ecological constraints on zooxanthellate corals

**DOI:** 10.64898/2026.03.18.712438

**Authors:** Kelly Gomez-Campo, M. Isabel Martinez-Rugerio, Miguel Ángel Gómez Reali, Raúl A. González-Pech, Michel Geovanni Santiago-Martínez, Kira E. Turnham, Todd C. LaJeunesse, Iliana B. Baums, Susana Enríquez, Roberto Iglesias-Prieto

## Abstract

Ecological models using light limitation to explain coral depth distribution have largely disregarded the energetic cost of sustaining photosynthetic activity. Here, we quantified photosystem II (PSII) turnover across a depth-simulated light gradient in a zooxanthellate coral, measuring PSII half-life, D1 protein abundance, and PSII-complex gene expression. Maximum photosynthetic capacity remained stable across irradiance levels while respiration rose and PSII turnover accelerated as a power law, imposing increasing ATP demand at the shallowest depths. Declining D1 protein abundance alongside stable transcript levels demonstrated that this escalating maintenance cost operates through post-transcriptional regulation. Consequently, a decreasing fraction of photosynthetic usable energy is available for translocation to the coral host at high irradiance, as the energy required for PSII repair increases. Integrating these physiological constraints into a bio-optical model revealed that the balance between photosynthetic capacity and its maintenance cost defines an optimal depth, the Photosynthetic Usable Energy Supply (*PUES*) maximum, where host energetic returns are maximized. This framework provides a mechanistic basis for understanding depth distributions in symbiotic corals and extends as a predictive tool for any photosynthetic organism operating under variable irradiance, including forecasting how environmental degradation contracts viable depth ranges.

## Introduction

Light supplies the energy that sustains coral reef ecosystems, but its role in shaping coral distribution cannot be explained by light availability alone. Variation in light quantity and spectral composition set physiological limits that structure community composition, species richness, and colony morphology across depth gradients (*1–6*). These ecological patterns emerge from fundamental biophysical constraints on how efficiently light energy is captured and converted into metabolic output. Corals are modular, sessile, mixotrophic organisms that host intracellular dinoflagellates in the family Symbiodiniaceae (*7*). Endosymbionts translocate a large fraction of their photosynthetically fixed carbon to the host, fueling basal metabolism and calcification (*8–10*). Light harvesting by the coral holobiont is determined by the emergent optical properties of symbionts, host tissue, and skeleton (*11*). Consequently, efficient photosynthesis requires coordinated adjustment across multiple levels of biological organization, from symbiont cell pigmentation and density, to host tissue optics, skeletal architecture, and whole colony morphology (*1*, *12–26*). These traits are integrated through photoacclimatory processes that optimize energy conversion under different light environments (*24*, *27–30*). Yet, the physiological limits through which photosynthesis constrains coral ecological niches across depth remain unresolved (*5*, *9*, *31*, *32*).

Depth gradients impose continuous selective pressures on coral–symbiont associations as sunlight simultaneously determines energy acquisition and physiological demands. If coral distribution were determined by solar energy alone, productivity and species richness would be expected to peak in shallow reefs and decline monotonically with depth as light attenuates. This pattern would reflect reduced growth rates and increased extinction risk under lower energy supply (*33*). However, for symbiotic corals, light availability by itself often fails to predict vertical patterns of biodiversity and performance (*3*). Instead, the amount of light that can be converted into biologically usable energy for the host better explains species richness and depth zonation in zooxanthellate corals (*5*). This distinction has relevant ecological implications. At intermediate depths, many corals maximize their net usable energy, expanding niche space and promoting greater species coexistence relative to both shallow and deep extremes. Under this framework, depth zonation cannot be simply considered a response to light attenuation with depth, but the outcome of selection on traits that determine how efficiently corals convert depth-specific available light into net energetic return. This is achieved through coordinated optimization of coral light capture, physiological adjustments of symbionts and host, and colony morphology. While lower depth limits are attributed to light limitation, upper depth limits are hypothesized to be constrained by the increasing energetic demands required to maintain performance under high irradiance.

Among the physiological processes that impose these energetic demands, protein damage and repair rank among the most energetically expensive cellular processes across living systems (*34–37*). However, their contribution to whole-organism energy budgets remains difficult to quantify. In photosynthetic organisms, this challenge is exemplified by the photosystem II (PSII) reaction center D1 protein, which undergoes rapid, light-dependent cycles of damage, degradation, and resynthesis (*38–43*). Replacement of photodamaged PSII reaction centers requires substantial ATP investment to support D1 synthesis, assembly, and regulation (*44–47*). In coral-associated Symbiodiniaceae, accelerated D1 turnover has been documented under both light- and thermal stress, consistent with a central role in sustaining photosynthetic function under stress conditions and maintaining the stability of the mutualism (*48–50*). Thus, empirical data on the energetic cost of sustaining this repair activity, and its consequences for energy partitioning within the symbiosis, are critical to determine energy utilization and delineate energetic boundaries in depth gradients.

We hypothesize that changes in PSII damage and repair across a light gradient affect the net energy that can be translocated from the symbiont to the coral host. Enhanced repair in high light environments sustains photosynthesis but requires more energy for ATP production and protein synthesis, reducing the fraction of photosynthetic energy available in support of host metabolism. Conversely, low light conditions limit energy production regardless of maintenance demands. These mechanisms predict a non-linear relationship between light availability and net autotrophic benefit to the coral host, providing an explanation for why abundant light in shallow reefs does not necessarily translate into enhanced coral growth or physiological performance (*5*).

To measure the effect of metabolic cost across high and low light conditions, we conducted a series of experiments to characterize coral photoacclimatory responses, quantify energetic demands of PSII damage-repair activity, and its impact on energy translocation to the coral host (**Fig. S1**). First, (1) we engineered a custom light system capable of simulating natural diurnal light cycles under controlled irradiance and spectral quality. This system minimized the confounding effects of ultra violet (UV) light (**Fig. S2**) and stabilized daily light integrals (DLI in mol quanta m⁻² day⁻¹) against the variability caused by cloud cover (*51*, *52*). Second, (2) coral fragments from the depth generalist *Orbicella faveolata* were acclimated to DLI treatments for four weeks, followed by a detailed characterization of its optical, structural, and physiological light phenotypes, including PSII half-life and turnover dynamics (**Table 1**). Third, (3) coral fragments were exposed to short-term shifts in DLI to evaluate repair capacity under light conditions that induce light limitation and light stress. Lastly, (4), we developed a bio-optical model integrating empirical results and theoretical considerations to predict shifts in photosynthetic usable energy available to the coral host across a depth gradient. This mechanistic framework provides a tool to predict the natural habitat and depth distribution of various coral taxa as well as quantify the impact of environmental changes and increase turbidity (diminished water quality) on the energetic cost to reef-building corals.

**Table 1.**
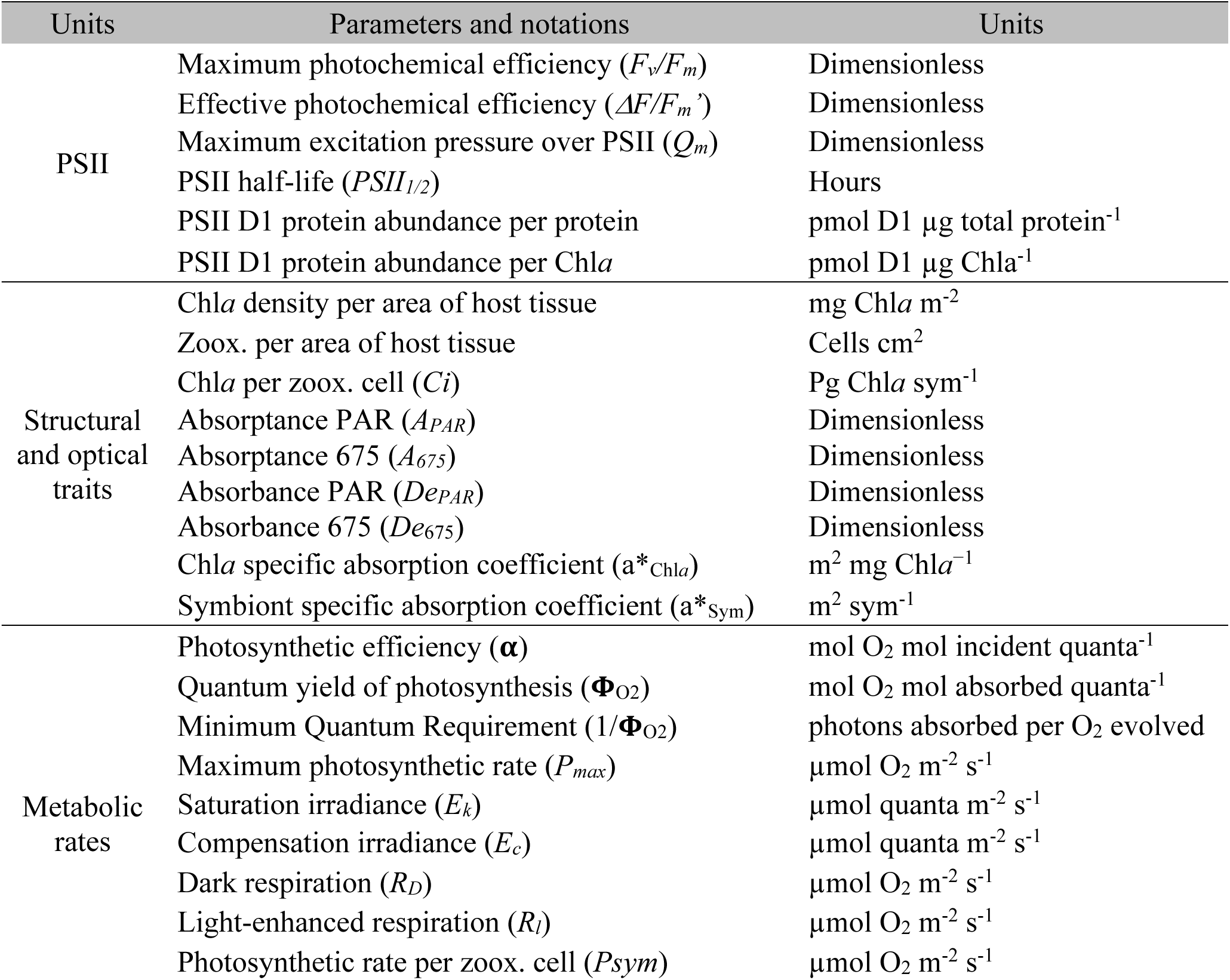
Table of terms. Descriptors used for phenotyping coral holobionts in the different levels of organization.

## Results

### Rapid photoacclimation across a realistic depth-dependent light gradient

Colonies in replicates were exposed to daily light integrals of 4 (DLI-4), 12 (DLI-12), 22 (DLI-22), and 32 (DLI-32) mol quanta m^-2^ day^-1^ (**Fig. 1A**, **Fig. S3A**), which simulates a depth gradient of 5–34 m (assuming a diffuse attenuation coefficient for downwelling irradiance, *K_d_*, of 0.07 m^-1^; typical for clear-water tropical reefs). PSII maximum photochemical efficiency (*F_v_/F_m_*) for experimental replicates (sample fragments) reached a steady state by day 10 (**Fig. S3B**), indicating that samples had acclimated to each of their respective daily doses of irradiance.

**Fig. 1.**
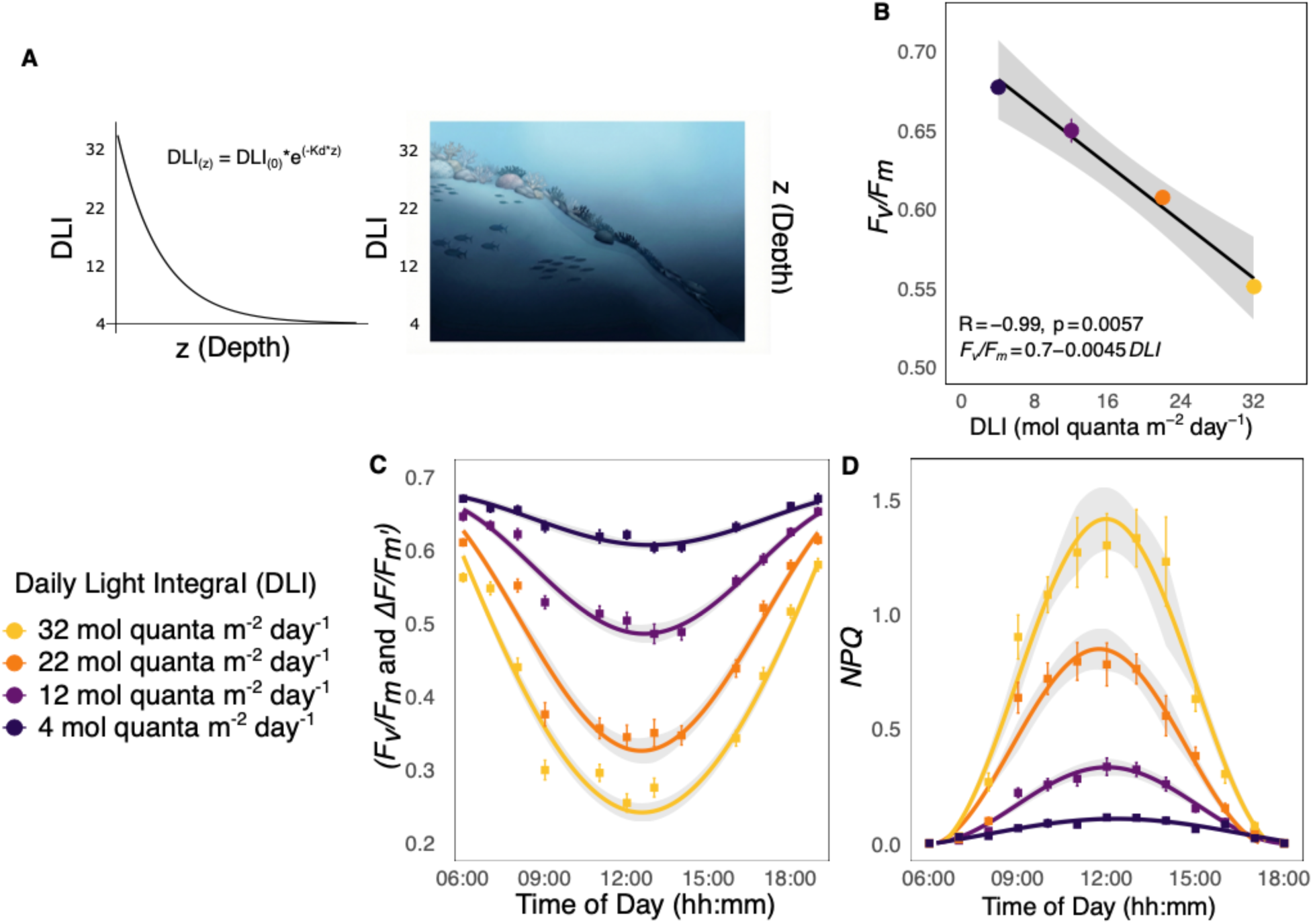
Phenotyping after acclimation to light conditions. (**A**) The general equation of light attenuation in seawater (DLI_(z)_ = DLI_(₀)_ × e ^(-Kd×z)^; Eq. 1) maps the four experimental DLI conditions to their corresponding depths (5–35 m, assuming *K_d_* = 0.07 m⁻¹), illustrated alongside a depth gradient. (**B**) Maximum photochemical efficiency (*F_v_/F_m_* at dusk) as a function of DLI after 4+ weeks of acclimation illustrating a linear trend. Adjustments in photoprotection shown as (**C)** diurnal changes in photochemical efficiency (maximum *F_v_/F_m_* at dawn/dusk, and effective *ΔF/F_m_*_’_ during daylight) and (**D**) non-photochemical quenching (*NPQ*) measured hourly across light conditions. Symbols represent mean ± SE (n = 16). Shaded areas show bootstrapped 95% confidence intervals.

### Diurnal photoprotection scaled with light exposure

Traits were treated as continuous functions of DLI rather than as discrete treatment means, characterizing phenotypic adjustments across the light gradient (**Fig. S4**). *F_v_/F_m_* was highest under DLI-4 and declined linearly with increasing light intensity (**Fig. 1B**). To quantify diurnal variation in PSII excitation pressure, we measured maximum pressure over PSII (*Qm*), which increased linearly corresponding to the transition from low to high irradiance (**Fig. S3C**). *Qm* was positively correlated with non-photochemical quenching (*NPQ*; r^2^ = 0.96), consistent with increases in photoprotective energy dissipation capacity as excitation pressure increased.

PSII photochemical efficiencies, maximum (*F_v_/F_m_*) and effective (*ΔF/F_m_^’^)*, measured every hour captured diurnal variation in PSII performance over the diurnal cycle. Differences in photochemical efficiency during the day were driven by light exposure, with a characteristic decline during the morning, a minimum at midday, and a reciprocal recovery in the afternoon (**Fig. 1C**). These diurnal dynamics reflected the balance among photoprotective mechanisms: induced dissipation as heat of the energy absorbed in excess (*NPQ*), induced PSII inactivation, and recovery (including *NPQ* relaxation) over the course of the day. The amplitude of diurnal oscillations increased proportionally with light exposure, reaching a maximum under the highest light treatment (DLI-32), which resulted in progressive increases in *NPQ* (**Fig. 1D**).

### Optical reorganization reduces bulk light absorption under high irradiance

Coral tissue pigmentation darkened or lightened in response to the corresponding light treatment, resulting in systematic changes in tissue optical properties (**Fig. 2A–B, Fig. S3D, Table S1**). From minimum to maximum exposure, Chlorophyll *a* (Chl*a*) content declined by ≈ 36% (112 ± 13 to 41 ± 4 mg Chl *a* m⁻²), while intact tissue absorbance peak at 675 nm (*De₆₇₅*) decreased by ≈ 11%. This corresponded to a decline in average absorptance in the PAR range (*A_PAR_*) of 6% from low to high light conditions. The reduction in Chl*a* pigment and slight reduction in bulk absorption capacity resulted in a non-linear increase in holobiont efficiency to absorb light (chlorophyll-specific absorption coefficient a*_Chl*a*_) (**Fig. 2B**).

**Fig. 2.**
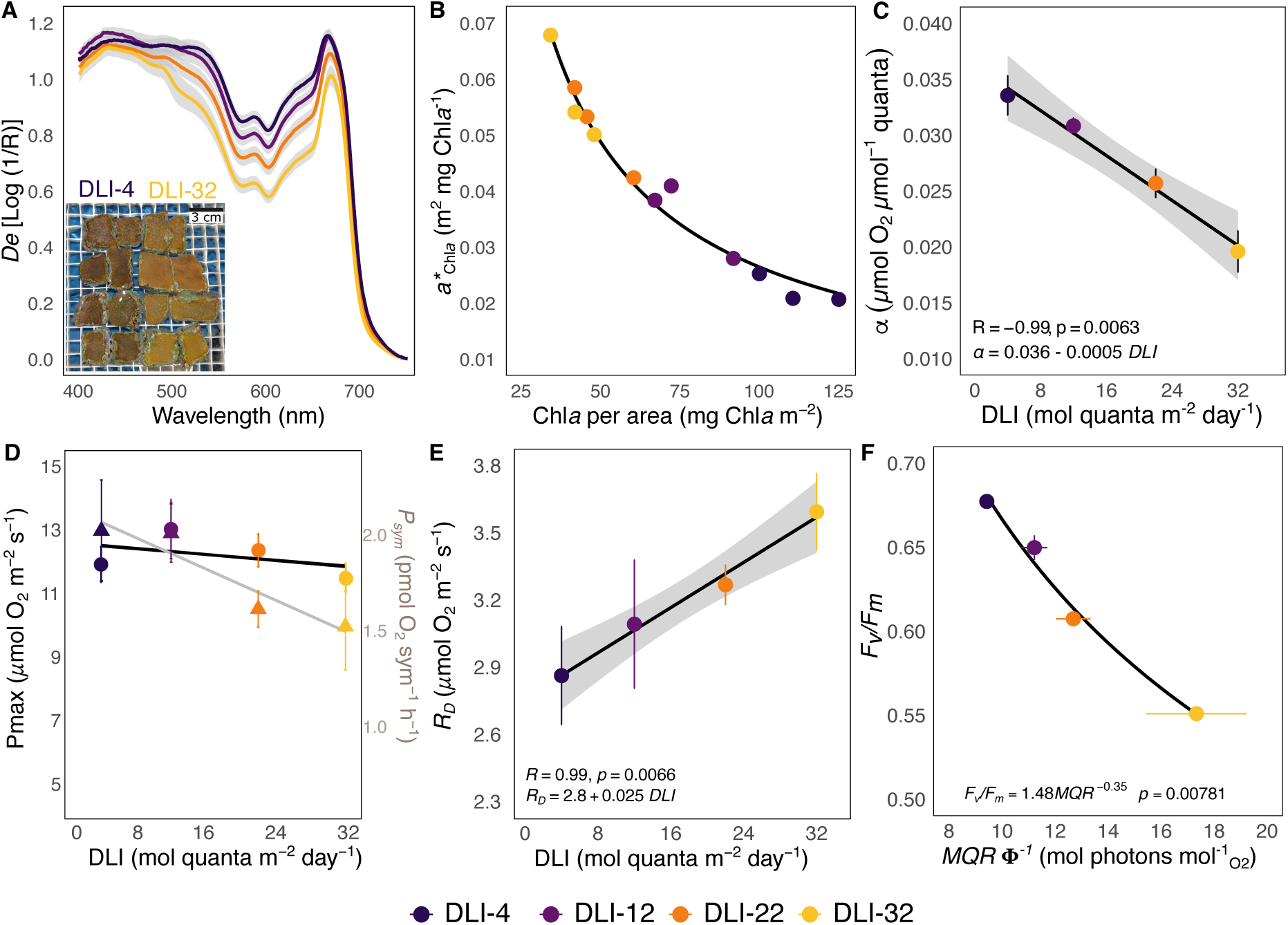
Optical reorganization and metabolic rates for light harvesting across light conditions. (**A**) Estimated absorbance (*De*) from reflectance measurements indicating reduced absorption under high light (n = 16) and (photo) qualitative changes in tissue pigmentation. (**B**) Pigment-specific absorption coefficient (a*\**_Chl*a*_) showing the non-linear increase in absorption efficiency in the coral tissue as pigment content declines, consistent with the package effect. (**C**) Photosynthetic efficiency (α), (**D**) maximum photosynthetic rate (*Pₘₐₓ*normalized to host area, and *P_sym_* normalized to symbiont cell) showing no significant changes in *P_max_* across light regimes (n = 8). (**E**) Dark respiration rates (*R_D_*) (n = 8), and (**F**) maximum photochemical efficiency of PSII (*F_v_/F_m_*) as a function of the minimum quantum requirement for photosynthesis (MQR, Φ⁻¹; mol photons mol⁻¹ O₂). Lower MQR values indicate higher photosynthetic efficiency or fewer photons required per molecule of O₂ evolved. DLI-4 corals achieved MQR values approaching the theoretical minimum (∼8 quanta O₂⁻¹), indicating near-maximal photosynthetic efficiency and delineating the lower boundary of photoacclimatory capacity for the symbiosis. With increasing irradiance (toward DLI-32), both *F_v_/F_m_* and quantum efficiency declined, reflecting the increasing energetic cost of sustaining photosynthesis under high light. Shaded areas show bootstrapped 95% confidence intervals; symbols are means ± SE.

### Photosynthetic capacity is maintained despite increasing respiratory demands

Metabolic rates were quantified using photosynthesis–irradiance (P/E) curves fitted with a hyperbolic tangent model, from which four photosynthetic parameters were extracted: photosynthetic efficiency (*α*), maximum photosynthetic rate (*P_max_*), saturation irradiance (*E_k_*), and compensation irradiance (*E_c_*) (*53*). Photosynthetic efficiency (α) increased as light exposure decreased, indicating enhanced efficiency under low irradiance conditions (**Fig. 2C**). In contrast, maximum photosynthetic rate (*P_max_*) remained stable across all light treatments at approximately 12 ± 0.33 μmol O₂ m⁻² s⁻¹ (**Fig. 2D**), indicating that photoacclimation preserved maximal photosynthetic capacity despite large differences in light exposure and pigmentation.

P/E curves allowed estimation of dark respiration rates (*Rd*), which increased linearly with light exposure (**Fig. 2E**), indicating a higher ATP demand in corals acclimated to higher light conditions. The phototrophic contribution to host energy requirements also increased with light exposure, as reflected by the higher daily integrated photosynthesis-to-respiration ratios (*54*) (**Fig. S3E**). The minimum quantum requirements of oxygenic photosynthesis (**Φ**^-1^_O2_) ranged from 10 to 17 quanta O₂⁻¹ across treatments (**Fig. 2F**) and the quantum efficiency of the photosynthetic process (**Φ**_O2_) from 0.1 to 0.059 mol O_2_ released per mol quanta absorbed. Both parameters revealed values close to the theoretical minimum of 8 quanta O₂⁻¹ for **Φ**^-1^_O2_ and the theoretical maximum of 0.125 for **Φ**_O2_ (*55*, *56*) for functional oxygenic photosynthesis.

### PSII functional half-life shortens sharply with increasing light exposure

Having quantified how absorbed light energy is dissipated as excitation pressure increases, and the resulting metabolic adjustments, we next examined processes regulating maintenance of functional PSII reaction centers under contrasting light conditions. First, to separate damage accumulation (before noon) from repair (before dusk), we inhibited *de novo* protein synthesis using chloramphenicol (CAP) (**Fig. 3A**). In control corals (−CAP), PSII activity declined through midday and recovered by the end of the light cycle as expected, whereas CAP-treated corals (+CAP) showed sustained loss of PSII activity consistent with inhibited repair.

**Fig. 3.**
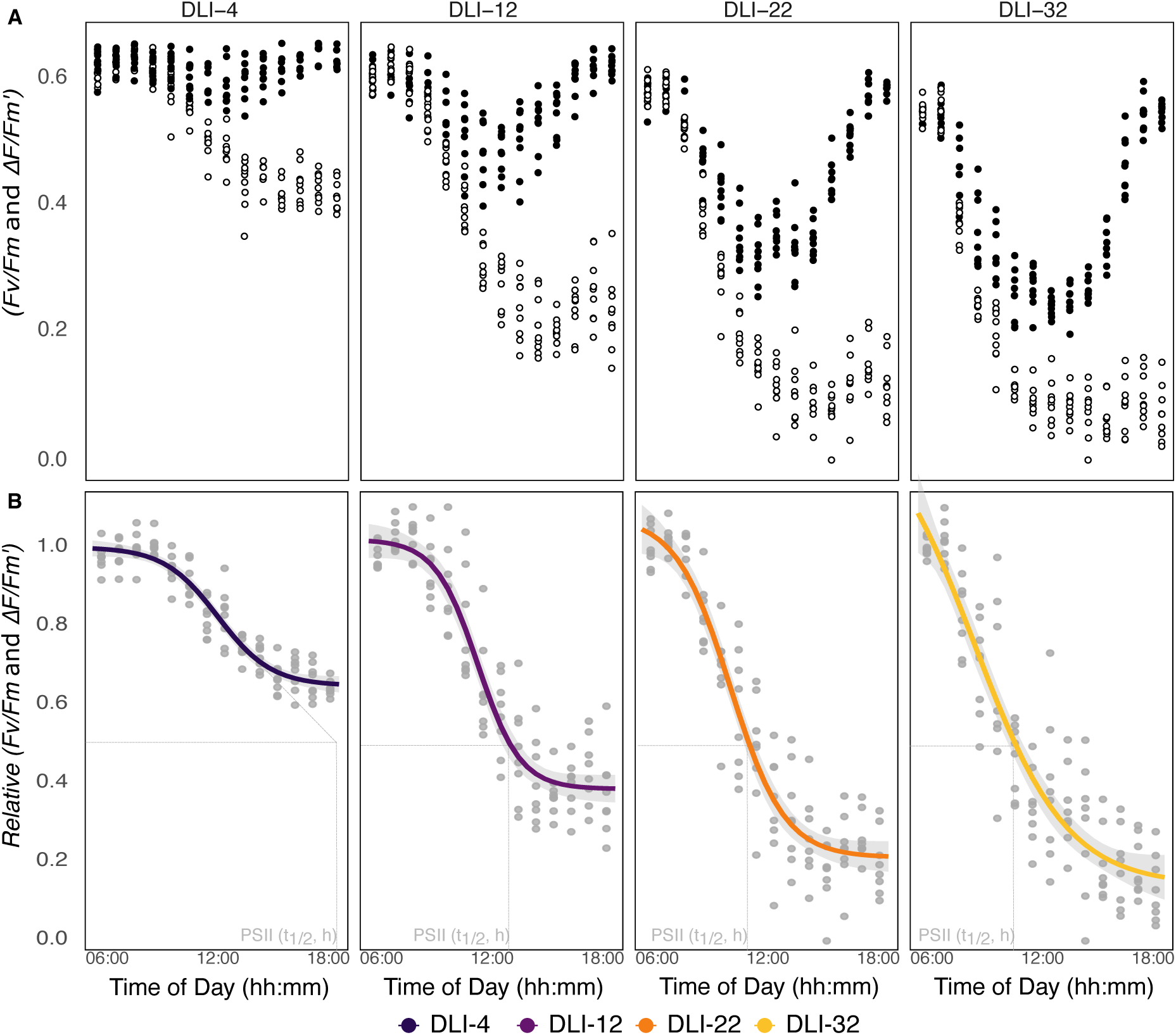
Diurnal PSII activity and PSII half-life. (**A**) PSII photochemical efficiency measured hourly across a 12 h photoperiod in the presence (+CAP; empty circles) and absence (−CAP; filled circles) of the protein synthesis inhibitor chloramphenicol. Blocking *de novo* protein synthesis revealed net PSII damage accumulation, while control samples showed oscillations with characteristic daily photodamage and repair cycles. (**B**) Relative photochemical efficiency (+CAP/-CAP) decay showing proportional rate of photodamage with increased light intensity, reflecting the rate of PSII inactivation in the absence of repair. Dashed lines illustrate the time of day when 50% of PSII reaction centers are photo-inactivated. In DLI-4 PSII t₁/₂ exceeded the 12 h photoperiod. Curves show four-parameter logistic fits (mean ± 95% CI).

This approach enabled estimation of PSII functional half-life (t₁⁄₂, h), defined as the number of daylight hours required to inactivate 50% of functional PSII reaction centers (**Fig. 3B**). PSII half-life declined exponentially from ∼13 h under low light to ∼3 h under high light exposure (**Fig. 4A**). This resulted in PSII turnover rates (τ_PSII_ day⁻¹) increasing significantly from low to high light exposure (**Fig. 4B**), revealing accelerated replacement of photodamaged reaction centers under high irradiance.

**Fig. 4.**
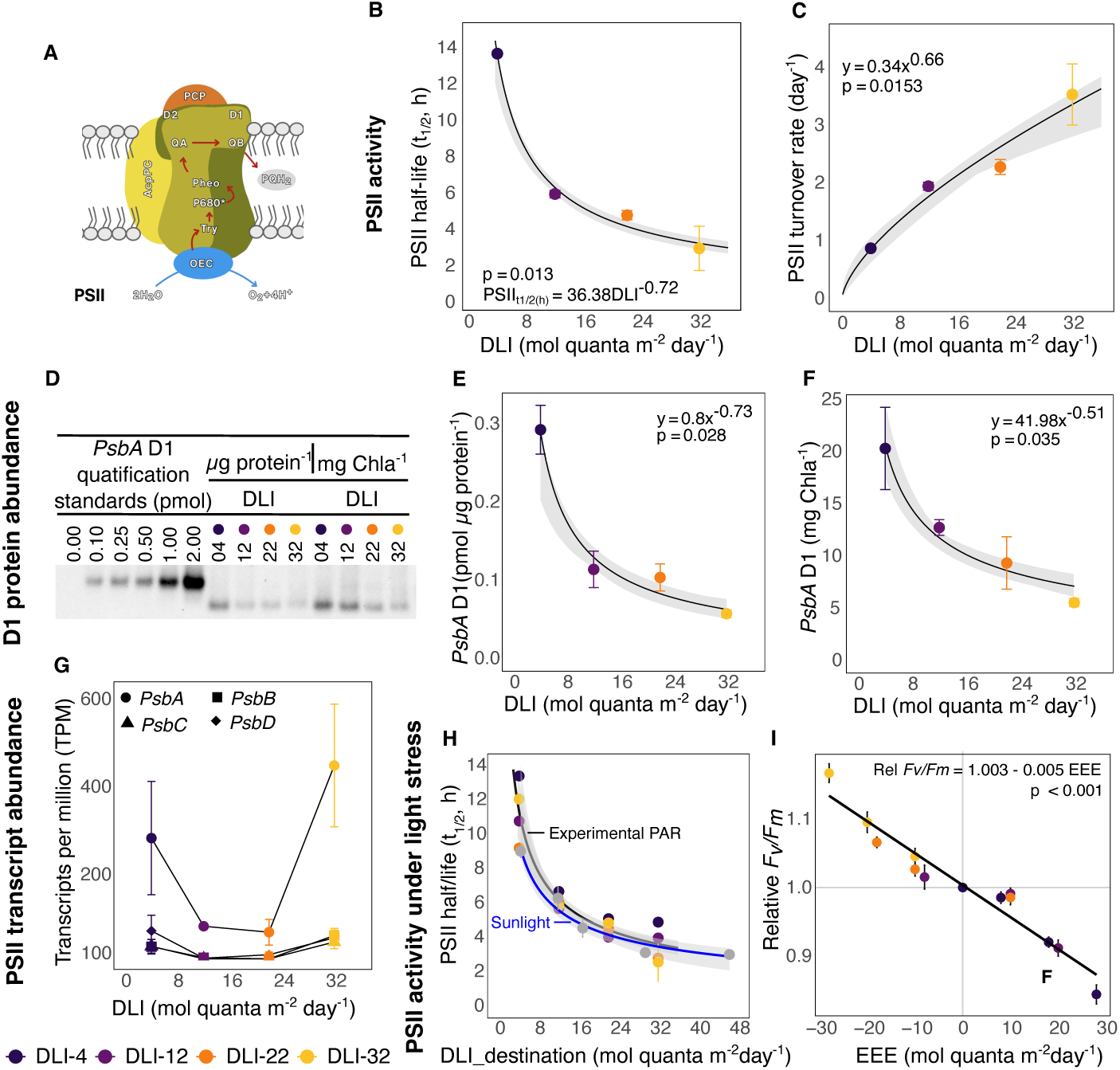
PSII regulation in zooxanthellate corals across light conditions. (**A**) PSII large pigment–protein complex with its core reaction center D1 (encoded by *psbA* gene). D1 provides key ligands to the oxygen-evolving complex, harbors the QB plastoquinone binding site, and is the primary target of photodamage. (**B**) Power law decay of PSII half-life (t₁/₂, h) with increasing DLI. (**C**) PSII turnover rate (day^-1^) accelerated significantly under higher DLI. (**D**) Quantitative immunoblot of the D1 protein, showing patterns across light conditions. Lanes include a negative control (lane1, 0.00 pmol), quantification standards (lanes2–6, 0.10-2.00 pmol), and samples loaded by total protein (lanes7–10, total protein^-1^) and Chl*a* content (11–14, mg Chl*a*^-1^). Quantification of D1 protein normalized to total protein (**E**; pmol D1 protein µg total protein⁻¹) and Chl*a* (**F**; pmol D1 protein mg Chl*a*⁻¹), showing a similar power law decay with light exposure (n = 3 immunoblots). (**G**) Transcriptional abundance (transcripts per million; TPM) of PSII complex genes (*psbA, psbB, psbC, psbD*). Transcript levels showed no systematic relationship with DLI, in contrast to the exponential decline in D1 protein abundance shown in panels E and F, indicating post-transcriptional regulation. (**H**) PSII half-life (t₁/₂, h) when acclimated corals (DLI) were changed to an alternative light condition (DLI-destination). Lines show a power law fit to experimental PAR (black line and colored circles; this study) and natural sunlight measurements (blue line and grey circles; data from (*5*)). Shaded areas show bootstrapped 95% confidence intervals; symbols are means ± SE. (**I**) Photodamage accumulation after the change in light condition. Results are shown as the relative change in *F_v_/F_m_* (*relative-F_v_/F_m_* = *F_v_/F_m_*-destination / *F_v_/F_m_*-acclimation) when coral fragments were exposed to *EEE* (*EEE* = DLI-destination - DLI-acclimation); the linear prediction is consistent with prior findings (*57*).

To assess molecular regulation underlying these dynamics, we quantified transcriptional and translational responses of major PSII reaction center proteins. Quantitative immunoblotting revealed significant differences in D1 protein abundance across the light conditions (**Fig. 4D–F**). D1 content declined sharply with increasing light exposure, mirroring the exponential decrease in PSII half-life and supporting regulation at the translational or post-translational level (**Fig. 4E**). Normalization of D1 content to chlorophyll *a* content yielded the same pattern (**Fig. 4F)**. In contrast, transcript abundance of the core components of the reaction center *psbA* (D1) and *psbD* (D2), and inner antenna proteins (that flank the D1/D2 heterodimer) *psbB* (CP47) and *psbC* (CP43), showed no systematic relationship with light treatment (**Fig. 4G**; **Fig. S5**).

### PSII half-life rapidly adjusts to acute changes in light exposure

To quantify short-term adjustments in PSII half-life under acute light stress, corals acclimated to a given DLI were transferred to the three alternative light conditions for a day, following the same CAP protein synthesis inhibition protocol (**Fig. S6**). PSII half-life responded rapidly to the new light environment and was independent of the prior acclimatory state, indicating that short-term regulation was driven by the destination light condition rather than the acclimation history (**Fig. 4H**).

To place these responses in the appropriate environmental context, we compared changes in half-life of PSII activity under our experimental PAR, and under natural sunlight, which also includes ultraviolet radiation (UV). In both cases, PSII half-life exhibited comparable exponential declines as a function of the destination light condition (**Fig. 4H**). Corals transferred from lower light treatments (DLI-4, DLI-12, DLI-22) to the highest light level (DLI-32) converged on a PSII half-life of approximately 3 h, matching that of colonies acclimated to DLI-32. These results demonstrate that PSII half-life can be rapidly adjusted within a single day across a wide range of light environments.

Despite this rapid adjustment, short-term regulation was insufficient to prevent the accumulation of PSII photodamage under high excitation pressure conditions. The magnitude of light stress, conveyed as the balance between PSII damage and repair, and quantified as *relative-F_v_/F_m_* changes (see methods) (*57*), declined with increasing excess excitation energy (EEE) (**Fig. 4I**). This indicated an increase in the magnitude of physiological light stress due to the rapid accumulation of damaged PSII.

### Energetic cost of photoacclimation constrains usable energy across depth

The bio-optical model developed here integrates theoretical considerations, empirical measurements, and equations describing phenotypic adjustments to predict the photosynthetic energy available for the coral host in depth-dependent light gradients (see methods). In a previous model implementation (*5*), community-averaged photophysiological responses were represented using a lumped-parameter approach, with photosynthetic and maintenance parameters optimized to reproduce observed biodiversity patterns across depth. Here, we incorporated light-dependent photoacclimatory profiles to explicitly predict photosynthetic energy allocation.

First, we computed DLI as a function of depth across a range of diffuse attenuation coefficients (*K_d_* = 0.04–0.40 m⁻¹), spanning from clear to turbid water conditions, to account for variation in underwater light environments, (**Eq. 1–2; Fig. S7**). Then, for each optical scenario, daily gross photosynthetic output (*DPg_(z)_*) was estimated from photosynthesis–irradiance curve parameters (**Eq. 3–4**), while depth-specific maintenance cost (*ECMpt_(z)_*) was derived from empirically constrained PSII turnover rates (**Eq. 5**). These components were computed to predict the photosynthetic usable energy supply (*PUES*) available to the coral host after accounting for symbiont metabolic demands (**Eq. 6**).

Across all optical scenarios, *PUES* exhibited a unimodal relationship with depth (**Fig. 5A**). In shallow waters, high irradiance required elevated maintenance cost to sustain photosynthetic rates. Consequently, an increasing fraction of photosynthetic energy was allocated to symbiont maintenance rather than host translocation. At greater depths, light limitation reduced carbon fixation, constraining energy availability despite lower PSII repair activity. Importantly, the depth at which *PUES* was maximized shifted systematically with *K_d_* (**Fig. 5B**). Consequently, clearer waters supported deeper energetic optima, whereas increased turbidity compressed the energetic window for coral distribution toward shallower depths. These results define an optimal bio-optical state for coral performance that emerges from the balance between photosynthetic output and maintenance cost, and whose position and extent depend on water-column optical properties.

**Fig. 5.**
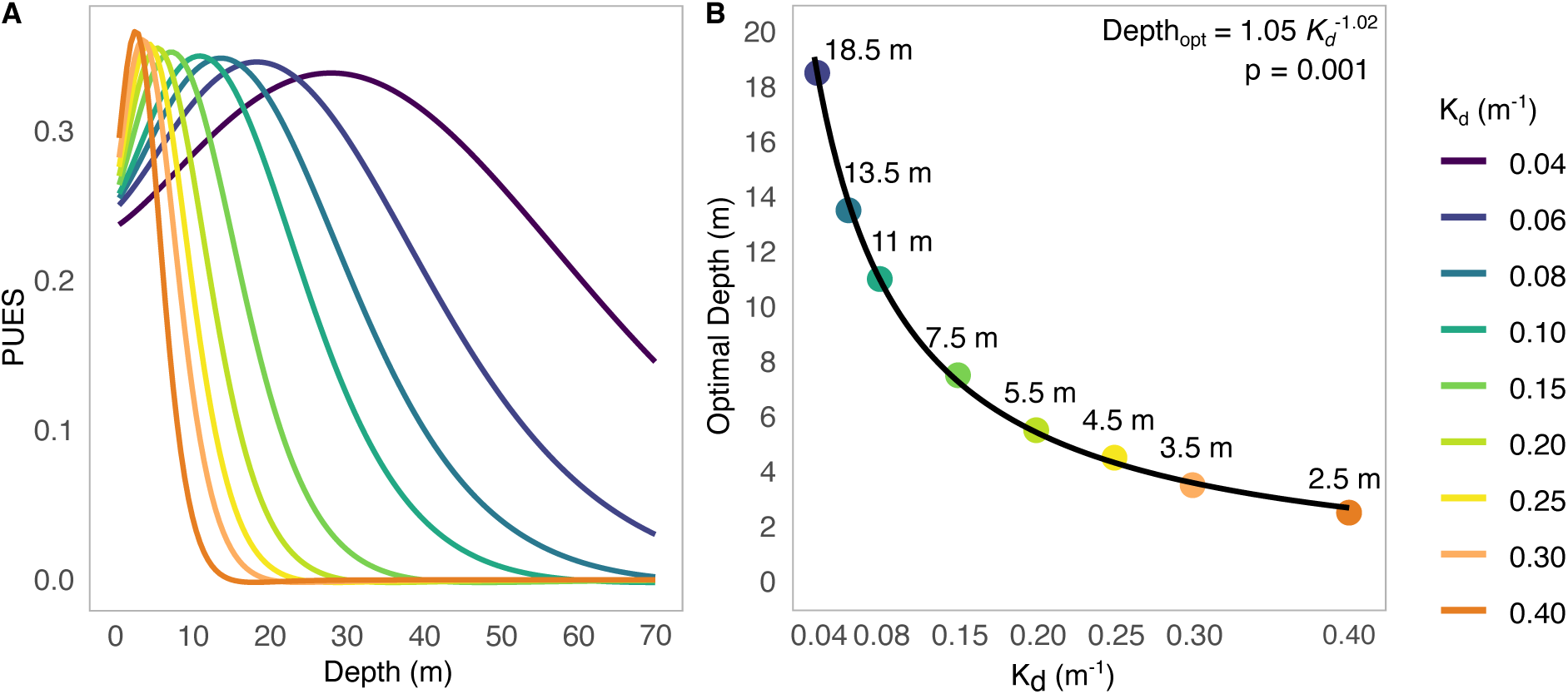
Bio-optical model of *photosynthetic usable energy supply* (*PUES*). (**A**) Modeled profiles of *PUES* across depth for a range of vertical attenuation coefficients of downwelling irradiance (*K_d_* = 0.04–0.40 m⁻¹), representing varying water column optical properties. *PUES* represents the photosynthetic energy available to the coral host after subtracting the depth-dependent energetic cost of PSII repair. Each curve represents a distinct *K_d_* value, color-coded from clear to turbid conditions. (**B**) Optimal depth (*Depth_opt_*) at which *PUES* is maximized, as a function of Kd. This bio-optical state represents the depth at which the balance between photosynthetic output and PSII repair cost is optimized, maximizing the energy potentially available for host metabolic demands.

## Discussion

This investigation provides a mechanistic basis for disentangling the apparent paradox between light availability, coral physiological performance, and vertical distribution of coral species by demonstrating that energetic constraints arise at both ends of the depth gradient. Clearly, carbon acquisition is limited by low irradiance. However, in shallow waters, high light intensity increases the metabolic cost of maintaining photosynthetic rates. Across a typical light gradient, maximum photosynthetic capacity remained stable (**Fig. 2D**) despite rising excitation pressure (**Fig. 1B, Fig. S3C**), enhanced photoprotective capacity (**Fig. 1C-D**), and elevated respiration (**Fig. 2E**). This homeostasis under high irradiance was sustained by accelerated PSII turnover and shortened reaction center half-lives (**Fig. 4**), imposing increasing ATP demand. Consequently, greater light availability (*i.e.* energy acquisition) did not directly translate into greater net energetic benefit to the host. Instead, a growing fraction of photosynthetic usable energy was retained by the symbiont for its cellular maintenance.

A bio-optical model based on these physiological dynamics predicted a unimodal relationship between light and usable energy supply (**Fig. 5A**), defining an optimal depth range bounded by the maintenance cost of living in shallow waters and carbon limitation at depth (**Fig. 5B**). Photosynthetic Usable Energy Supply *(PUES)* was previously conceptualized in the context of Species–Energy Theory (*33*) to explain patterns of coral biodiversity (*5*); here we provide the physiological basis for these distributions by incorporating the metabolic cost of sustaining photosynthesis under a wide range of light conditions. Classic carbon-budget studies show that zooxanthellae translocate a large proportion of their net daily fixed carbon to the coral host, with the fraction of host respiratory demand varying substantially with species, depth, and light regime (*8*, *58*). Translocated products such as glycerol, glucose, and other metabolites that vary among Symbiodiniaceae lineages (*7*, *59–63*) support host metabolism, growth, and calcification, whereas a smaller fraction is retained for symbiont cell growth (*58*, *64*). Our results demonstrate that this translocation is predictable. As irradiance increases, ATP-intensive photoprotective processes, particularly rapid PSII repair via D1 turnover, reduce the fraction of organic carbon available for host use.

The cost of maintaining cellular function is rarely included in photosynthesis or primary productivity models, which traditionally rely on photosynthesis:respiration ratios and/or Chl*a* to identify conditions of positive carbon balance (*5*, *65–67*). By coupling depth-dependent photosynthetic output with associated maintenance cost, *PUES* reframes coral energetics by emphasizing photosynthetic usable energy for the host. The resulting bio-optical model provides a tool to translate physiological limits to the ecological distribution of photosynthetic organisms, which in the case of zooxanthellate corals, involves colony growth and morphology as well as gamete production and the timing of reproduction. Additionally, it offers a framework for predicting how environmental stressors contract depth distributions by necessitating greater investment in metabolic maintenance.

The stability of *P_max_* across the range of light exposure reveals the central energetic problem: photoacclimation in the coral holobiont preserves photosynthetic capacity despite large changes in light availability. Disentangling the mechanisms requires examining each component of the acclimatory response – photoprotection, optical adjustment, metabolic adjustments, and reaction center maintenance – and what each reveals about the limits of metabolic flexibility.

The first and energetically cheapest line of defense against excess irradiance is thermal dissipation via *NPQ*. The tight coupling between *Q_m_* and *NPQ* (r² = 0.96) reflects the dynamic photoprotective equilibrium, driven by xanthophyll cycling, that scales continuously with excitation pressure across diel cycles and light conditions (*1*, *16*, *68–71*), consistent with prior field and laboratory observations (*16*, *70*, *72–75*). Critically, however, *NPQ* alone cannot prevent photodamage when excitation pressure exceeds the capacity for thermal energy dissipation. The declining *F_v_/F_m_* with increasing irradiance marks precisely this transition, which is the point where the *NPQ* buffer is insufficient and repair via D1 turnover becomes the dominant maintenance strategy, with an escalating ATP cost (*47*, *88*, *89*).

The optical organization of the coral host and its zooxanthellae operate under strict acclimatory limits at both ends of the depth gradient. Coral skeleton and tissue optics have evolved for light amplification rather than attenuation (*11*, *24*). Despite a nearly threefold decline in Chl*a* content across light conditions, overall absorptance remained effectively constant, with total energy absorbed preserved regardless of pigment reduction (**Table S1**). Rather than reducing total light absorption, pigment loss increased chlorophyll-specific absorption efficiency (a*_Chl*a*_) as the holobiont approached its high-light acclimatory limit (*11*, *76*, *77*). At this boundary, the capacity for further pigment reduction is exhausted, leaving the symbiosis more vulnerable to additional stressors such as thermal stress (*78*), which is consistent with the higher occurrence of bleaching in high-irradiance environments. The same optical architecture that enables colonization of light-limited deep reefs (*27*, *29*) simultaneously constrains the upper irradiance tolerance of the symbiosis, as it prevents meaningful reduction of absorbed energy through pigment adjustment alone. Thus, in shallow reef environments, when this optical limit is reached and repair capacity is overwhelmed, selection favors colony morphologies with structural self-shading, *i.e.* branching, foliose, and massive forms with strong within-colony light attenuation (*27*, *30*, *79*).

The stability of *P_max_* across the range of light exposure marks a physiological constraint on carbon fixation, not merely a correlate of photoacclimation. Additional irradiance above saturation does not generate additional fixed carbon, hence it can only be dissipated as heat or directed toward repair. Evidence for this limit emerges from both ends of the light gradient. Low light phenotypes approached the theoretical maximum for PSII photochemical efficiency in dinoflagellates *in hospite* (*F_v_/F_m_* ≈ 0.7) and the theoretical minimum quantum requirement for oxygen evolution (∼10 quanta O₂⁻¹; **Fig. 2F**) (*55*, *56*), marking the lower boundary of photoacclimatory capacity (*1*, *54*). On the other hand, high light phenotypes, while more efficient at absorbing light (higher a*_Chla_), were proportionally less efficient at converting it into fixed carbon (lower *α* and *Φ_max_*). The symbiosis thus operates across a bounded photoacclimatory range between light-limited in deep water and light-saturated in shallow water, with *P_max_* as the stable outcome. This reveals that coral photoacclimation operates through modulation of light absorption, heat dissipation, and protein turnover rather than by regulating photosynthetic capacity. Such decoupling suggests that photosynthetic rates are constrained by downstream processes, such as inorganic carbon diffusion or alternative electron sinks within the symbiosome (*80–83*) and, more critically, demonstrates that maintaining photosynthetic performance under high light requires progressively greater metabolic support without proportional energetic return. The primary source of this increasing cost is PSII turnover and D1 protein dynamics.

Protein turnover is among the most energetically expensive cellular processes (*34–37*). In higher plants, thylakoid protein turnover accounts for 13–38% of total ATP consumption, with D1 replacement representing a major fraction of this budget (*37*). Energy for this repair cycle is supplied through respiration, linking PSII maintenance directly to metabolic demand. By preventing protein synthesis in the symbiont, we decoupled PSII damage from repair and quantified changes in PSII half-life across experimental light conditions. PSII half-life declined significantly with increasing irradiance and closely paralleled reductions in D1 protein abundance. Accordingly, corals acclimated to high light conditions maintained maximum photosynthetic capacity via faster replacement of photodamaged reaction centers and lower D1 pools. Similar relationships between D1 abundance, PSII half-life, and light history have been documented in higher plants, including how rapid D1 turnover accompanies elevated metabolic demand under high light exposure (*37*, *84–86*). The elevated dark respiration (*R_d_*) found under high light conditions is therefore consistent with increased ATP requirements for D1 synthesis and PSII reassembly (*36*, *44*, *87*). ATP availability is known to constrain *de novo* D1 synthesis and full PSII repair (*47*, *88*, *89*), suggesting that ATP supply becomes the limiting currency sustaining photosynthetic function under light stress.

In this context, D1 turnover operates as a regulated sink for excess excitation, preventing over-reduction of the electron transport chain and protecting PSI under acceptor-side limitations (*45*, *86*). This hierarchy reflects structural cost optimization within the photosynthetic apparatus. Because antenna complexes are far more abundant (∼7.6 × 10⁶ cell⁻¹) and energetically more expensive than D1 protein (∼1.0 × 10⁵ cell⁻¹) (*90*), it makes selective turnover of PSII reaction centers a comparatively economical strategy. In fact, regulation of reactive oxygen species within the photosynthetic apparatus shows that while chlorophyll triplet states in antenna complexes are efficiently quenched by carotenoids, spatial constraints within the PSII reaction center prevent such protection, allowing singlet oxygen to target D1 directly and requiring rapid turnover as a controlled safeguard (*40*, *91*, *92*). D1 protein, then, becomes the unavoidable target of oxidative damage, making its rapid turnover a necessary mechanism to preserve photosynthetic integrity.

This feature of D1 damage and its costly turnover has direct ecological implications. Earlier work has primarily focused on how PSII repair impairment under thermal stress leads to coral bleaching. When repair fails to match damage rates, D1 protein abundance drops below the threshold needed to sustain photosynthetic activity, resulting in pigment loss, symbiont degradation, and ultimately, expulsion or symbiont digestion by the host (*48*, *93–96*). In our experiments, even in the absence of heat stress, high light and PAR alone shortened PSII half-life from ∼13 h to ∼3 h (**Fig. 4B**) and reduced D1 abundance by 80% (**Fig. 4E**), imposing substantial ATP demand. Thus, the upper tolerance limits of the coral holobiont are defined not solely by photodamage, but by the energetic capacity to sustain PSII repair. This constraint is always active in high light conditions, revealing the high susceptibility of shallow water corals to any additional environmental stress. We confirmed the environmental regulation of this mechanism by comparing PSII half-life under experimental PAR (400–700 nm) and natural sunlight (PAR + UV) (**Fig. 4H**). No significant differences in PSII half-life were found, consistent with PAR being the primary driver of photoinactivation and daily damage accumulation. This result challenges the assumption that UV is the dominant driver of PSII photodamage in shallow-water corals (*83*, *97*).

Ultimately, PSII repair is a fundamental energetic ‘bottleneck’ that constrains coral holobiont performance at high irradiance. Sustaining photosynthesis under light stress requires continuous ATP investment in protein repair and replacement, thereby diverting resources from growth, reproduction, and calcification. In this way, energetic maintenance cost of photosynthesis sets the upper limits of energy acquisition by reef corals and their stress tolerance to episodes of acute environmental change.

A bio-optical model incorporating Photosynthetic Usable Energy Supply provides a mechanistic scaling of coral bioenergetics to explain their ecological patterns of depth and latitude distribution (**Fig. 5A**). The model predicts the fraction of photosynthetically derived energy that is available for host metabolic requirements, including maintenance, growth, reproduction, and ecological interactions, and thus, defines fundamental ecological niches of the coral holobiont. In the bioenergetic framework, the *PUES* bio-optical model, specifically *PUES* optimum (*Depth_opt_*), defines the depth at which photosynthetic supply and maintenance demand are optimally balanced, maximizing usable energy translocation while minimizing the cost of sustaining photosynthetic integrity (**Fig. 5B**). The *Depth_opt_* position and extent of coral colonization depend on water column optical properties. When exploring a range of light attenuation coefficients (*K_d_*), spanning clear to turbid conditions, our analysis revealed that increased light attenuation shifts *Depth_opt_* toward shallower waters and compresses the energetic envelope. Therefore, environmental degradation, including progressive ocean darkening (*i.e*., increased turbidity), drives a compression of the viable depth range for coral colonization and growth.

The development of adaptations to better meet the energetic needs for living at various depths and water conditions carries significant evolutionary consequences. Intermediate depths, where energy use is maximized without excessive physiological costs, support multiple photophysiological strategies and species coexistence. In contrast, the higher energetic costs of protein maintenance in shallow water or extremes of carbon limitation at greater depth, ultimately restrict possible trait combinations channeling ecological success through specialization under strong selection pressure. Such patterns align with general Species-Energy Theory and with emerging genomic evidence that closely related coral lineages repeatedly diverge across narrow depth gradients, with adaptation to distinct light conditions linked to environmental sensing and reproductive timing (*98–100*).

## Materials and Methods

### PAR LED Light System

We developed an experimental light system by engineering custom-made lamps and software able to simulate diurnal light cycles. Natural light in the tanks was absent throughout the experiments. The illumination system relied on 100W light-emitting diodes (LED)-lamps that enabled automation of the diurnal cycles (**Fig. S3A**). The four light treatments (Daily Light Integral, DLI) were: 4 mol quanta m^-2^ day^-1^ (DLI-4, peak of irradiance 200 μmol quanta m^-2^ s^-1^); 12 mol quanta m^-2^ day^-1^ (DLI-12; peak of irradiance 450 μmol quanta m^-2^ s^-1^); 22 mol quanta m^-2^ day^-1^ (DLI-22; peak of irradiance 750 μmol quanta m^-2^ s^-1^); and 32 mol quanta m^-2^ day^-1^ (DLI-32; peak of irradiance 1100 μmol quanta m^-2^ s^-1^).

Three parameters were prioritized for the control of each light condition: light intensity, light quality and time. We built each LED-lamp with a BC Series High CRI COB-LED-400L-100W-Unit (YujiLEDS®) which best covered the sunlight spectrum in the range of the Photosynthetic Active Radiation (PAR). LEDs were operated in continuous mode with a multifunction I/O card (USB-6001, National Instruments Corp., USA) to avoid potential artifacts on photosynthesis and fluorescence responses related to the “flashing effect” of commercial LEDs at different pulsating frequencies (*101*). Light intensity was controlled with a custom-made software in LabVIEW (Laboratory Virtual Instrument Engineering Workbench, National Instruments Corp., USA). To eliminate natural sunlight inside experimental tanks, we placed LED-lamps inside black boxes that fully covered each tank (**Fig. S1**).

Diurnal light exposure (mol quanta m^-2^ day^-1^) was described by integrating the per second change in irradiance (µmol quanta m^-2^ s^-1^) over a day as Daily Light Integral (DLI). Light exposure as an integral has proven to be a useful tool for assessing irradiance accumulated when the response of interest is growth or acclimation (*102*). DLIs conditions captured a wide range of coral vertical distribution in clear waters (5 to 35 m depth for a downwelling light attenuation coefficient, *K_d_*, of 0.07 m^-1^). Irradiance at depth depends on the optical properties of the water column given by the *Kd* at any reef location. Irradiance was calibrated and continuously monitored with the quantum sensor LI-1400 (LI-COR, USA).

Light spectra also play an important role in photosynthesis performance. The effective activation of the repair cycle of photodamaged PSII has its own action spectrum. Studies have shown that UV-radiation damages PSII more effectively than visible light (*97*), and that the reported action spectra are rather dissimilar in the range between 400 and 700 nm. We used a sun-like spectrum and UV was excluded by design (**Fig. S2**). We were able to effectively induce PSII damage repair cycles in the simulated DLIs treatments.

### Acclimation to Daily Light Integrals

We sampled 64 fragments from 4 *Orbicella faveolata* colonies (each a different genet) between 4-5 m depth under similar light conditions in La Bocana, Puerto Morelos National Park. Experimental coral fragments (∼5-9 cm^2^) were set for a 2-week healing period after sampling by placing them on PVC-deployed structures located at 5 m depth near the sampling area. Corals were then transported to the National Autonomous University of Mexico (UNAM) mesocosm facility, where each colony was equally distributed in four outdoor tanks (shaded natural sunlight to 16 ± 3 mol quanta m^2^ day^-1^) equipped with running natural seawater (n = 16 fragments per tank). Water temperature and flow were maintained at 28 ± 1°C and 1100 L h^-1^ using commercial aquaria chillers and heaters. Water temperature in the tanks was continuously monitored with Hobo data loggers (Onset Computer Corporation, MA, USA). Corals were acclimated to the mesocosm for two weeks before LED installation, followed by 4+ weeks under experimental DLI conditions before phenotyping.

We monitored daily PSII maximum photochemical efficiency (*F_v_/F_m_*) at dusk and effective photochemical efficiency (Δ*F/F_m_’*) at noon, with a pulse-amplitude modulated fluorometer (Diving-PAM; Heinz Walz, Effeltrich). Maximum excitation pressure over PSII (*Q_m_* = 1 - [(Δ*F/F_m_’* at noon) / (*F_v_/F_m_* at dusk)]) was also estimated to monitor the moderate light stress induced during natural and daily photo-acclimation, and the adjustment to a stable photo-acclimatory condition (*1*).

### Phenotyping in the range distribution of zooxanthellate corals

Following the 4+ weeks of acclimation to the experimental light gradient, we measured a suite of traits to characterize coral phenotypes (**Table 1**). Chl*a* fluorescence measurements were recorded as photochemical efficiency (*F_v_/F_m_* and Δ*F/F_m_’*), and non-photochemical quenching was calculated as *NPQ* = [(*F_m_-F_m’_*)/*F_m_*].

To describe changes in coral pigmentation, we estimated Chl*a* and Chl*c* contents. Briefly, an air gun and filtered sea water (FSW) were used to obtain tissue slurries subsequently homogenized at low temperature (Tissue-Tearor Homogenizer BioSpec Inc, USA) and centrifuged. The resulting pellet was re-suspended in filtered seawater and preserved for pigment concentration (extracted with acetone/dimethyl sulfoxide 95:5 vol/vol) and Symbiodiniaceae cell density (counted in a haemocytometer after the addition of 200 μL of iodine preservation solution). Chl*a* and Chl*c* concentrations were estimated spectrophotometrically (3 reads per sample) with a modular spectrometer (Flame-T-UV-VIS, Ocean Optics Inc., USA) using the equations described by Jeffrey and Humphrey (*103*).

Light absorption spectra were determined on intact *in vivo* coral tissues from reflectance (*R*) measurements. *R* was used to estimate Absorbance (*De*) [*De* = log (1/*R*)] and Absorptance (*A*), assuming that Transmittance through the coral skeleton was negligible (*104*). *A* was estimated for the PAR range (400 and 700 nm; *A_PAR_* = 1 – *R_PAR_*); and at the 675 peak of Chl*a* (*A_675_* = 1 – *R_675_*).

Specific-absorption coefficients (a*) were normalized to Chl*a* content (a*_Chl*a*_ ; m^-2^ mg Pigm^-1^) as [a*_Chl*a*_= (*De*_675_/*ρ*)·ln(10), where *De_675_* = log(1/*R_675_*)] (*11*, *77*, *105*). Likewise, a* was normalized to symbiont cell density (a*_Sym_; m^-2^ sym^-1^) as [a*_sym_ = (*De_675_/*sym *ρ*) · ln(10)]. These descriptors estimate the holobiont efficiency to absorb light (a*_Chl*a*_) and symbiont specific light absorption efficiency (*a*\*_sym_) (*78*).

To describe changes in coral metabolic rates, we used photosynthesis / irradiance curves (P/E) fitted to a hyperbolic tangent model (**Fig. S4A**). Briefly, coral fragments were incubated using a laboratory-made water-jacketed respirometer (*76*) with filtered sea water at a constant temperature (28 °C) and constant water flow generated by continuous agitation from magnetic stirrers, which ensure that the boundary layer around coral surfaces is broken. One chamber was measured as a blank (no coral fragment) to correct for background activity in the water. A LED-system was designed to enable automation of light increments every 10 min. Oxygen evolution was measured continuously with a fiber-optic oxygen meter system (FireSting, Pyroscience) to estimate the rate in the 10 min range of: dark (pre-illumination), light (10 light increments), dark (post-illumination) for a total period of the 2 h incubation. Photosynthetic efficiency (*α*), compensation irradiance (*E*_c_), saturation irradiance (*E*_k_), respiration rates in the dark (*R*_D_), light-enhanced respiration (R_L_), and maximum photosynthetic rates (*P*_max_), were estimated from the PE curves (*106*, *107*). Each curve reflects the photosynthetic potential of the light phenotype. To estimate the energy absorbed in excess, we calculated the hours of light exposure above *E_k_* (number of hours in *P_max_*; H_sat_ (*108*)) and the magnitude of energy absorbed under these conditions (mol quanta m^-2^ day^-1^).

Using absorptance values determined above [*A* = 1- *R*], we calculated the maximum quantum yield of photosynthesis (**Φ**_O2_; mol oxygen evolved per mol quanta absorbed^-1^) and its inverse, the minimum quantum requirements of photosynthesis [(**Φ**^-1^_O2_; = 1/(α * *A_PAR_*); quanta required to release an O_2_ molecule^-1^)] (*106*, *107*). **Φ**_O2_ was estimated from α, the slope of the linear regression of photosynthesis in the sub-saturating region of the P/E curve; corrected for the light absorbed (*A*) in our experimental setting [(photosynthetic utilizable radiation (PUR) absorbed (*A_PUR_*)] (*76*, *106*, *107*). To estimate the integrative index of the holobiont’s metabolism, we calculated daily integrated photosynthesis-to-respiration ratios [*Pg (E)* day^-1^: *R (E)* day^-1^].

### Algal symbiont identification

*Orbicella faveolata* hosts distinct species of algal symbionts in this region (*e.g.*, *Symbiodinium* sp. A3, *Breviolum faviinorum*, *Breviolum meandrinum*, *Cladocopium* sp. C3, *Cladocopium* sp. C7, and *Durusdinium trenchii*), with the possibility of multiple species simultaneously inhabiting a same colony (*109–111*). To account for this functional diversity, symbiont identity was determined using two complementary approaches.

(1) One cm^2^ coral fragments were used to extract genomic DNA using a modified Promega Wizard protocol (Promega Madison, WI) (*112*, *113*). The large ribosomal subunit (*LSU* rDNA) was amplified using *28S* forward (5’-CCCGCTGAATTTAAGCATATAAGTAAGCGG-3’) and *28S* reverse (5’-GTTAGACTCCTTGGTCCGTGTTTCAAGA-3’) (*114*) and an annealing temperature of 66 °C. Amplifications were Sanger sequenced using the Big-Dye Terminator 3.1 Cycle Sequencing Kit (ThermoFisher Scientific, Waltham, MA) and analyzed on the Applied Biosciences Sequencer (Applied Biosciences, Foster City, CA) at the Penn State University Genomics Core Facility. Sequences were checked and edited in Geneious v11.0.5 (Biomatters Ltd, Auckland, NZ) and were queried against the NCBI nucleotide database using BLASTn (National Center for Biotechnology Information).

(2) We separately aligned raw RNA-Seq reads (see below) from each sample against reference transcriptomes using bwa v0.7.18 (*115*); mapping rates were determined using SAMStat v2.2.3. The reference transcriptomes encompassed eight datasets from the most closely related Symbiodiniaceae that *Orbicella* associates with: *Symbiodinium* sp. A3 (*116*), *Breviolum faviinorum* (*116*), *Breviolum* sp. B5 (*116*), *Cladocopium proliferum* (*117*), *Cladocopium* sp. C1 MI (*118*), *Cladocopium* sp. C1 SM (*118*), and *Durusdinium trenchii* (*117*, *118*). Symbionts present in each sample were determined by comparing *LSU* rDNA and mapping results (**Fig. S5A**).

### PSII half-life and turnover rates under acclimatory conditions

After the 4-week acclimation, the PSII damage and repair cycle was determined through monitoring the variation of *F_v_/F_m_* and *τιF/Fm’* in 12 h diurnal cycles. We tracked the accumulation of photo-inactivated PSII and the accumulation of photo-repaired PSII. The damage accumulation phase runs from dawn to midday, and repair from midday to dusk.

To quantify the effect of light acclimation on the photodamage and repair of the PSII active pool, we inhibited the PSII repair process. In each acclimatory condition, fragments were separated into two subgroups: control (n = 8, no inhibitor -CAP) and the inhibitory treatment (n = 8, treated with D1 protein synthesis inhibitor chloramphenicol; +CAP). Subgroups were placed separately in 1 L plastic containers with internal seawater circulation (FSW) and positioned within tanks (water baths). Water was exchanged in the containers every 2 h for both treatments (FSW+CAP) and controls (FSW-CAP). PSII photochemical efficiency was measured every hour, and samples were snap frozen after dusk for further analyses. This protocol was repeated for every light condition after 4+ weeks of acclimation. To estimate PSII half-life (t_1/2_, h) photochemical efficiency treatment-to-control ratios [+CAP / -CAP] were curve fitted using a logistic sigmoidal function and expressed for each light condition as the number of hours required to inactivate 50% of PSII (t_1/2_, h).

### Abundance of PSII D1 protein

Algal pellet assigned for protein analyses was flash frozen in liquid nitrogen and stored at -80 °C. We quantified D1 protein content in algal symbionts isolated from three coral fragments for each light regime. Each immunoblotting assay was performed multiple times to load: (lane 1) a negative control, (lanes 2–6) 0.10-2.00 pmol *psbA*/D1 standard, (lanes 7–10) one sample from each light regime to be normalized by total protein (μg protein^-1^), and (lanes 11–14) one sample from each light regime to be normalized by Chl *a* (mg Chl *a*^-1^). We performed quantitative immunoblotting following established protocols (*93*, *94*, *96*, *119*, *120*). Briefly, proteins were extracted using protein extraction buffer PEB (Agrisera AS08 300) supplemented with Pefabloc® SC (Sigma-Aldrich 11429868001) to inhibit proteolysis. Samples were subjected to three cycles of sonication-to-liquid N_2_ to break cell wall and avoid over-heating. Protein concentrations were determined using The Better Bradford assay with a standard curve. Equal amounts of total protein were mixed with Bolt™ LDS sample buffer (Invitrogen B0007) and Bolt™ Sample Reducing Agent (ThermoFisher B0004), denatured, and separated on NuPAGE™ 4–12% Bis-Tris gels (Invitrogen NP0329BOX) alongside PageRuler™ Plus pre-stained protein ladder (ThermoScientific 26619). Following electrophoresis, proteins were transferred to PVDF membranes (Invitrogen LC2005) using Bolt™ transfer buffer (Invitrogen BT0006) and blocked with ECL™ Advance Blocking Reagent (GE Healthcare GERPN418). Membranes were probed with a rabbit polyclonal anti-PsbA (D1) antibody (Agrisera AS05 084), followed by HRP-conjugated chicken anti-rabbit secondary antibody (Agrisera AS10 833). Membranes were stained post-transfer with SimplyBlue™ SafeStain (Invitrogen LC6060). D1 protein quantification was achieved by co-loading known amounts of PsbA*/*D1 standard (Agrisera AS01 016S), imaging on a chemiluminescent imager (ChemiDoc BIO-RAD), and pixel-quantifying in ImageJ.

### Expression of PSII-complex genes

Expression of PSII genes from algal symbionts was assessed following an RNA-Seq approach. Total RNA was extracted from 16 fragments, 4 untreated fragments (-CAP) per light exposure. Each sample was homogenized in TRIzol reagent (Ambion, Life Technology) before centrifugation with chloroform for 15 min at 12,000 *g* at 4 °C. The aqueous phase was then isolated and cleaned using Qiagen RNeasy Mini kit (Qiagen), as per the manufacturer’s protocol with an additional on-column DNase treatment using RNase-Free DNase Set (Qiagen). To maximize concentration of eluted RNA, the same 35 μL of RNAse-free water was twice passed through the spin column for the final isolation step. Concentration and purity were pre-screened via Qubit spectrometer and quality assessed (RNA integrity number; RIN > 7) with Agilent 2100 Bioanalyzer (Agilent Technologies). Purified RNA was sent to Biomarker Technologies (BMK) GmbH where concentration and purity were re-analyzed (Qubit® dsDNA BR Assay Kit on a Qubit® 2.0. Fluorometer). RNA libraries for 2 × 150 bp paired-end sequencing were non-stranded, prepared using Hieff NGS Ultima Dual-mode mRNA Library Prep Kit for Illumina (Yeasen) model:13533ES96. Samples were run in platform Illumina NovaSeq X.

Raw reads were assessed for quality using FastQC v0.12.1 (github.com/s-andrews/FastQC) and results visualized using MultiQC v1.28 (*121*), followed by filtering and trimming using Trimmomatic v0.39 (*122*). Pseudoalignments of the processed read pairs and quantification of expression were done using the quant function of kallisto v0.51.1 (*123*) in two manners: against all reference transcriptomes combined into a single database and against separate transcriptomes according to the symbiont type identified per sample. For the combined database, all reference transcriptomes were concatenated removing redundant sequences using cd-hit-est v4.8.1 (*124*) at a sequence similarity threshold of 0.99. The transcripts per million (TPM) output by kallisto were used as measure of gene expression level. Genes of the core complex of PSII were identified by a BLASTn v2.16.0 (*125*) search of Symbiodiniaceae GenBank records (JN557844 for *psbA*, HG515018 for *psbB*, HG515019 for *psbC*, and HG515020 for *psbD*) as queries against the reference transcriptomes with an e-value ≤ 10^-5^. The BLAST hit with the highest mean expression across samples per genus was selected as the representative transcript for each gene.

### PSII half-life under light-stress conditions

To quantify the effect of light stress on the PSII photodamage and repair cycle, and the variation of PSII half-life (t_1/2_, h), we replicated the experimental protocol to inhibit the PSII repair process. However, here the 4-weeks acclimated corals were switched (new coral fragments for each experiment) to an alternative light exposure (*i.e.*, DLI-4_acclimated_ switched to DLI-12_destination_, DLI-22_destination_ and DLI-32 _destination_) for one diurnal cycle. The experimental protocol was repeated for each light exposure for a total of 12 experiments: DLI-12_acclimated_ switched to DLI-4_destination_, DLI-22_destination_ and DLI-32_destination_; DLI-22_acclimated_ switched to DLI-4_destination_, DLI-12_destination_ and DLI-32_destination_; and DLI-32_acclimated_ switched to DLI-4_destination_, DLI-12_destination_ and DLI-22_destination_.

The severity of light stress was quantified with the metric Excess Excitation Energy *EEE* (mol quanta m^-2^ day^-1^) as follows (*57*): (1) Conditions where *EEE* = 0 (*EEE_0_*) indicate that corals are well acclimated (*F_v_/F_m_* is steady day-to-day), no net light stress is induced, hence no photodamage is accumulated. (2) Conditions where *EEE* ≠ 0 (*EEE*_≠*0*_) indicate corals were switched to a different light condition and may experience light stress. *EEE* was quantified as the difference between DLI in the switched condition (DLI_destination_) and DLI acclimated (DLI_acclimated_). In the case that the switched condition resulted in lower light exposure, *EEE* shows negative values. The magnitude of the physiological stress was measured as the ratio between *F_v_/F_m_* in the switched condition and *F_v_/F_m_* acclimated, which results in the parameter *relative-F_v_/F_m_*.

PSII half-life (t_1/2_, h) was estimated following the same protocol described above for PSII half-life under acclimatory conditions as the number of hours required to inactivate 50% of the active PSII pool.

### Modeling the energetic cost of photo-acclimation

To estimate how the light environment shapes the energetic balance of the coral-algal symbiosis, we developed a bio-optical model that integrates theoretical light attenuation in the water column, gross photosynthetic output, and the energetic costs of PSII turnover. The analysis focused on the estimation of the potentially maximum photosynthetic output of the symbionts after subtracting the costs of maintenance of photosynthetic activity through PSII repair, which can be translocatable to the host. Other metabolic costs required by the maintenance and/or growth of algae and host were not considered into the model due to the difficulties of parameterization of the large diverse of biological processes potentially involved. Light availability at depth was expressed as daily light integrals (DLI) and computed from surface irradiance. Different light attenuation coefficients (*K_d_*) were used. Depth-dependent photosynthetic output was then estimated using a photosynthesis-irradiance relationship parameterized with function-valued phenotyping derived from experimental profiles. To capture the costs of maintaining photosynthetic function under different light regimes, we incorporated PSII turnover dynamics by linking PSII lifetime (t_1/2_, h) and its cost, maximum photochemical efficiency (*F_v_/F_m_*), and repair potential to DLI. The resulting framework yields the Photosynthetic Usable Energy Supply (*PUES*) by the symbionts to the host, a metric that represents the potentially maximum energy available for the coral host after subtracting the light-dependent costs of photosynthesis maintenance. Considerations and computations underlying the model proceeded as follows:

### Theoretical and empirical considerations

As light is the primary factor that regulates primary production and the photoacclimatory response of photosynthetic organisms, we first estimated the amount of light available at a given depth (*126*):

#### 1. Solar irradiance as a function of depth

The intensity of a parallel-beam radiation of a particular wavelength decreases along its path in pure water, obeying the Lambert Law. The general equation of light attenuation in seawater is (*126*):

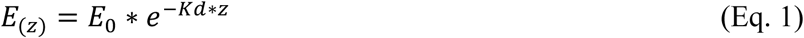

Where *E_(z)_* is light intensity at depth *z* (here irradiance value for Photosynthetic Active Radiation, PAR, as UVR was not considered), *E_0_* is the PAR just beneath the water surface (when *z* = 0) and *Kd* (m^−1^) is the vertical attenuation coefficient of downwelling PAR irradiance.

#### 2. Changes in daily light integral (DLI) with depth: Light exposure

Irradiance at any point on the Earth’s surface is determined by solar elevation. This rises during the day from zero at dawn to its maximum value at noon, and then diminishes in a precisely symmetrical manner to zero at dusk under no cloud cover. Light exposure integrates the variation in irradiance per second and m^2^ over time. The integrated quantum unit for light exposure at a certain depth and day is mol quanta m^-2^ day^-1^, which describes the amount of quanta received by m^-2^ during a 24 h period (*126*):

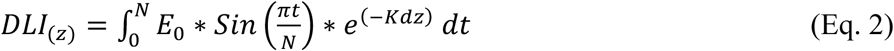

The parameter *DLI_(z)_* is then the daily light integral at depth, *E_0_* is the superficial irradiance at solar noon, *N* is the daylength (in seconds) and *t* is time along a day (seconds).

#### 3. Photosynthesis as a function of DLI

The response of photosynthetic organisms to available light (from the light saturation curve), depends on two fundamental parameters in our model: the slope α at very low light availability (carbon fixed/oxygen evolved per unit of available light), and the asymptotic maximum production *P_max_* (the rate of carbon fixation/oxygen evolved under saturating light) (*127*):

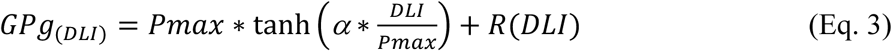

Where *GPg_(DLI)_* is the rate of gross photosynthesis measured at a given quantum irradiance, *DLI*; *P_max_* is the maximal photosynthetic capacity; α is the is the photosynthetic efficiency; and *R_(DLI_*_)_ is the light dependent variation of net coral respiration. As indicated previously, we used here the direct values of *R_L_* and *R_D_* measured for each coral phenotype, instead a single and general function for *Orbicella faveolata*, maintaining the same criteria: *R_L_* was used for the period of the day at photosynthesis saturation (*P_max_*), *R_D_* in darkness or very low light levels, and the average for the sub-saturating region, according to each acclimatory condition.

#### 4. Photosynthetic output as a function of depth

To estimate the photosynthetic output Δ*Pg(z)* at a given *z*, we needed to integrate daily photosynthesis at *z* (modified from (*127*)):

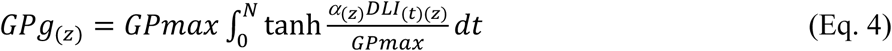

#### 5. Photosynthetic usable energy for coral maintenance

We tracked the relationship between photosynthetic and maintenance at depth and incorporated the energy cost of photoacclimation *PUES(z)* as the cost of protein turnover rate at depth. Energetic cost of 1 turnover (0 if no energy cost is involved in protein turnover) (*37*, *54*):

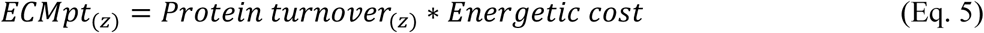

Photosynthetic usable energy supply for coral at depth z was expressed by integrating equations 4 and 5:

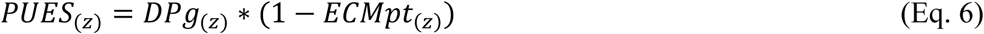

##### Function-valued phenotyping included in the model, where *z* is depth and DLI daily light integral

1. *Photosynthetic parameters as a function of DLI_(z)_:*
  a. Gross maximum photosynthetic rate *GPmax* = 15 μmol O_2_ m^-2^ s^-1^
  b. Saturation irradiance *Ek* as a function of *DLI_(z)_* (mol quanta m^-2^ day^-1^)

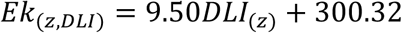
  c. Photosynthetic efficiency α as a function of *DLI_(z)_*

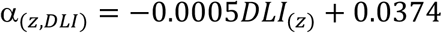
2. *Protein turnover rate as a function of DLI_(z)_:*
  a. PSII-half-time (hours) as a function of *DLI* (mol quanta m^-2^ day^-1^):

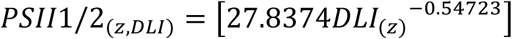
  b. Relative maximum photochemical efficiency (*Fv/Fm*):

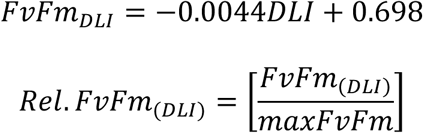
  c. Repair potential (dimensionless):

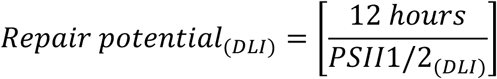
  d. Protein turnover rate:

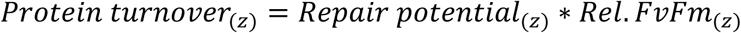

## Acknowledgments

We thank UNAM and ICMyL for staff support, facility assistance, SAMMO for data, S Enriquez for laboratory space and mesocosm facility. V Avila-Magana, CT Galindo-Martinez, NS Locatelli, CC Osborne for assistance in laboratory work. T López-Londono for assistance in modeling. VA Gómez-Chavez for Figure S1. Collection permit No. PPF/DGOPA-020/2018 and CITES permit No. MX96989, MX102782 to RIP and SE.

## Author contributions

Conceptualization: RIP, KGC.

Data curation: KGC.

Formal analysis: KGC and RIP.

Funding acquisition: RIP, SE, TCL, IBB.

Investigation and methodology: KGC performed the experiments analyze and integrated data. IMR contributed to fieldwork, phenotypic descriptions and light systems. MAGR engineered the light system. RGP to transcriptomic computational work and interpretations. MGS to translational laboratory work and interpretations. KET to algal identification laboratory work and interpretations. TLC and IBB to analysis and interpretations.

Project administration: RIP.

Resources: RIP, SE, TCL, IBB, MGSM.

Software: MAGR.

Supervision: RIP.

Validation: KGC, RIP.

Visualization: KGC, RIP.

Writing - original draft: KGC, RIP.

Writing - review & editing: KGC, IMR, MAGR, RGP, MGSM, KET, TCL, IBB, SE, RIP.

## Data and materials availability

Data is available in the main text or the supplementary materials. All detailed data, custom codes used in the analyses will be available for purposes of reproducing or extending the analyses in GitHub. All data generated by this study will be deposited to the National Center for Biotechnology Information (NCBI).

## References

1. R. Iglesias-Prieto, V. H. Beltrán, T. C. LaJeunesse, H. Reyes-Bonilla, P. E. Thomé, Different algal symbionts explain the vertical distribution of dominant reef corals in the eastern Pacific. Proceedings. Biological sciences / The Royal Society 271, 1757–1763 (2004).

2. G. S. E. Chow, Y. K. S. Chan, S. S. Jain, D. Huang, Light limitation selects for depth generalists in urbanised reef coral communities. Marine Environmental Research 147, 101–112 (2019).

3. T. E. Roberts, S. A. Keith, C. Rahbek, T. C. L. Bridge, M. J. Caley, A. H. Baird, Testing biodiversity theory using species richness of reef-building corals across a depth gradient. Biol Lett 15, 20190493 (2019).

4. R. Tamir, G. Eyal, N. Kramer, J. H. Laverick, Y. Loya, Light environment drives the shallow-to-mesophotic coral community transition. Ecosphere 10, e02839 (2019).

5. T. López-Londoño, K. Gómez-Campo, X. Hernández-Pech, S. Enríquez, R. Iglesias-Prieto, Photosynthetic usable energy explains vertical patterns of biodiversity in zooxanthellate corals. Sci Rep 12, 20821 (2022).

6. G. Pérez-Rosales, M. Pichon, H. Rouzé, S. Villéger, G. Torda, P. Bongaerts, J. Carlot, U. T. P. Consortium, V. Parravicini, L. Hédouin, Mesophotic coral ecosystems of French Polynesia are hotspots of alpha and beta generic diversity for scleractinian assemblages. Diversity and Distributions 28, 1391–1403 (2022).

7. T. C. LaJeunesse, J. E. Parkinson, P. W. Gabrielson, H. J. Jeong, J. D. Reimer, C. R. Voolstra, S. R. Santos, Systematic Revision of Symbiodiniaceae Highlights the Antiquity and Diversity of Coral Endosymbionts. Current Biology 28, 2570–2580.e6 (2018).

8. L. Muscatine, L. R. Mccloskey, R. E. Marian, Estimating the daily contribution of carbon from zooxanthellae to coral animal respiration. Limnology and Oceanography 26, 601–611 (1981).

9. P. G. Falkowski, Z. Dubinsky, L. Muscatine, J. W. Porter, Light and the Bioenergetics of a Symbiotic Coral. BioScience 34, 705–709 (1984).

10. J. P. Gattuso, D. Allemand, M. Frankignoulle, Photosynthesis and calcification at cellular, organismal and community levels in coral reefs: A review on interactions and control by carbonate chemistry. American Zoologist 39, 160–183 (1999).

11. S. Enríquez, E. R. Méndez, R. Iglesias-Prieto, S. Enrı, E. R. Me, Multiple scattering on coral skeletons enhances light absorption by symbiotic algae. Limnology and Oceanography 50, 1025–1032 (2005).

12. J. W. Porter, L. Muscatine, Z. Dubinsky, P. G. Falkowski, Primary Production and Photoadaptation in Light- and Shade-Adapted Colonies of the Symbiotic Coral, Stylophora pistillata. Proceedings of the Royal Society of London. Series B, Biological Sciences 222, 161–180 (1984).

13. R. K. Trench, Microalgal-Invertebrate symbioses : a Review. Endocytobiosis & Cell Research 9, 135–175 (1993).

14. R. Iglesias-Prieto, R. K. Trench, Acclimation and adaptation to irradiance in symbiotic dinoflagellates. II. Response of chlorophyll-protein complexes to different photon-flux densities. Marine Biology 130, 23–33 (1997).

15. W. K. Fitt, F. K. McFarland, M. E. Warner, G. C. Chilcoat, Seasonal patterns of tissue biomass and densities of symbiotic dinoflagellates in reef corals and relation to coral bleaching. Limnology and Oceanography 45, 677–685 (2000).

16. M. Y. Gorbunov, Z. S. Kolber, M. P. Lesser, P. G. Falkowski, Photosynthesis and photoprotection in symbiotic corals. Association for the Sciences of Limnology and Oceanography 46, 75–85 (2001).

17. E. A. Titlyanov, T. V. Titlyanova, K. Yamazato, R. van Woesik, Photo-acclimation dynamics of the coral *Stylophora pistillata* to low and extremely low light. Journal of Experimental Marine Biology and Ecology 263, 211–225 (2001).

18. K. R. N. Anthony, O. Hoegh-Guldberg, Kinetics of Photoacclimation in Corals. Oecologia 134, 23–31 (2003).

19. P. R. Frade, P. Bongaerts, A. J. S. Winkelhagen, L. Tonk, R. P. M. Bak, In situ photobiology of corals over large depth ranges: A multivariate analysis on the roles of environment, host, and algal symbiont. Limnology and Oceanography 53, 2711–2723 (2008).

20. S. J. Hennige, D. J. Smith, R. Perkins, M. Consalvey, D. M. Paterson, D. J. Suggett, Photoacclimation, growth and distribution of massive coral species in clear and turbid waters. Marine Ecology Progress Series 369, 77–88 (2008).

21. P. A. Todd, Morphological plasticity in scleractinian corals. Biological Reviews 83, 315–337 (2008).

22. M. P. Lesser, S. Marc, S. Michael, O. Michiko, R. D. Gates, G. Andrea, Photoacclimatization by the coral Montastraea cavernosa in the mesophotic zone: Light, food, and genetics. Ecology 91, 990–1003 (2010).

23. M. S. Roth, The engine of the reef: Photobiology of the coral-algal symbiosis. Frontiers in Microbiology 5 (2014).

24. S. Enríquez, E. R. Méndez, O. Hoegh-Guldberg, R. Iglesias-Prieto, Key functional role of the optical properties of coral skeletons in coral ecology and evolution. Proceedings of the Royal Society B: Biological Sciences 284, 20161667 (2017).

25. K. E. Lohr, E. F. Camp, U. Kuzhiumparambil, A. Lutz, W. Leggat, J. T. Patterson, D. J. Suggett, Resolving coral photoacclimation dynamics through coupled photophysiological and metabolomic profiling. J Exp Biol 222, jeb195982 (2019).

26. A. Malik, S. Einbinder, S. Martinez, D. Tchernov, S. Haviv, R. Almuly, P. Zaslansky, I. Polishchuk, B. Pokroy, J. Stolarski, T. Mass, Molecular and skeletal fingerprints of scleractinian coral biomineralization: From the sea surface to mesophotic depths. Acta Biomaterialia 120, 263–276 (2021).

27. M. O. Hoogenboom, S. R. Connolly, K. R. N. Anthony, Interactions Between Morphological and Physiological Plasticity Optimize Energy Acquisition in Corals. Ecology 89, 1144–1154 (2008).

28. Y. X. Ow, P. A. Todd, Light-induced morphological plasticity in the scleractinian coral Goniastrea pectinata and its functional significance. Coral Reefs 29, 797–808 (2010).

29. N. Kramer, R. Tamir, O. Ben-Zvi, S. L. Jacques, Y. Loya, D. Wangpraseurt, Efficient light-harvesting of mesophotic corals is facilitated by coral optical traits. Functional Ecology 36, 406–418 (2022).

30. K. Gomez-Campo, R. Sanchez, I. Martinez-Rugerio, X. Yang, T. Maher, C. C. Osborne, S. Enriquez, I. B. Baums, S. A. Mackenzie, R. Iglesias-Prieto, Phenotypic plasticity for improved light harvesting, in tandem with methylome repatterning in reef-building corals. Molecular Ecology 33, e17246 (2024).

31. P. Hallock, Algal symbiosis: A mathematical analysis. Mar. Biol. 62, 249–255 (1981).

32. M. O. Hoogenboom, K. R. N. Anthony, S. R. Connolly, Energetic cost of photoinhibition in corals. Marine Ecology Progress Series 313, 1–12 (2006).

33. D. H. Wright, Species-Energy Theory: An Extension of Species-Area Theory. Oikos 41, 496–506 (1983).

34. R. De Visser, C. J. T. Spitters, T. J. Bouma, “Energy cost of protein turnover: theorectical calculation and experimental estimation from regression of respiration on protein concentration of full-grown leaves” in Molecular, Biochemical and Physiological Aspects of Plant Respiration (1992), pp. 493–508.

35. T. J. Bouma, R. De Visser, J. H. J. A. Janssen, M. J. De Kock, P. H. Van Leeuwen, H. Lambers, Respiratory energy requirements and rate of protein turnover in vivo determined by the use of an inhibitor of protein synthesis and a probe to assess its effect. Physiologia Plantarum 92, 585–594 (1994).

36. A. Quigg, J. Beardall, Protein turnover in relation to maintenance metabolism at low photon flux in two marine microalgae. Plant, Cell and Environment 26, 693–703 (2003).

37. L. Li, C. J. Nelson, J. Trösch, I. Castleden, S. Huang, A. H. Millar, Protein degradation rate in Arabidopsis thaliana leaf growth and development. Plant Cell 29, 207–228 (2017).

38. G. Schuster, R. Timberg, I. Ohad, Turnover of thylakoid photosystem II proteins during photoinhibition of Chlamydomonas reinhardtii. European Journal of Biochemistry 177, 403–410 (1988).

39. E. M. Aro, S. McCaffery, J. M. Anderson, Photoinhibition and D1 Protein Degradation in Peas Acclimated to Different Growth Irradiances. Plant physiology 103, 835–843 (1993).

40. E.-M. Aro, I. Virgin, B. Andersson, Photoinhibition of Photosystem II. Inactivation, protein damage and turnover. Biochimica et Biophysica Acta (BBA) - Bioenergetics 1143, 113–134 (1993).

41. S. Järvi, M. Suorsa, E. M. Aro, Photosystem II repair in plant chloroplasts--Regulation, assisting proteins and shared components with photosystem II biogenesis. Biochimica et biophysica acta 1847, 900–909 (2015).

42. P. Chotewutmontri, A. Barkan, Dynamics of Chloroplast Translation during Chloroplast Differentiation in Maize. PLOS Genetics 12, e1006106 (2016).

43. L. Li, E. M. Aro, A. H. Millar, Mechanisms of Photodamage and Protein Turnover in Photoinhibition. Trends in Plant Science 23, 667–676 (2018).

44. K. Miyata, K. Noguchi, I. Terashima, Cost and benefit of the repair of photodamaged photosystem II in spinach leaves: Roles of acclimation to growth light. Photosynthesis Research 113, 165–180 (2012).

45. M. Tikkanen, N. R. Mekala, E. M. Aro, Photosystem II photoinhibition-repair cycle protects Photosystem i from irreversible damage. Biochimica et Biophysica Acta - Bioenergetics 1837, 210–215 (2014).

46. S. Järvi, J. Isojärvi, S. Kangasjärvi, J. Salojärvi, F. Mamedov, M. Suorsa, E.-M. Aro, Photosystem II Repair and Plant Immunity: Lessons Learned from Arabidopsis Mutant Lacking the THYLAKOID LUMEN PROTEIN 18.3. Front. Plant Sci. 7 (2016).

47. N. Murata, Y. Nishiyama, ATP is a driving force in the repair of photosystem II during photoinhibition. Plant, Cell & Environment 41, 285–299 (2018).

48. S. Takahashi, T. Nakamura, M. Sakamizu, R. Van Woesik, H. Yamasaki, Repair nachinery of symbiotic photosynthesis as the primary target of heat stress for reef-building corals. Plant and Cell Physiology 45, 251–255 (2004).

49. R. Hill, S. Takahashi, Photosystem II recovery in the presence and absence of chloroplast protein repair in the symbionts of corals exposed to bleaching conditions. Coral Reefs 33, 1101–1111 (2014).

50. S. Roberty, D. Fransolet, P. Cardol, J.-C. Plumier, F. Franck, Imbalance between oxygen photoreduction and antioxidant capacities in Symbiodinium cells exposed to combined heat and high light stress. Coral Reefs 34, 1063–1073 (2015).

51. K. R. N. Anthony, P. V. Ridd, A. R. Orpin, P. Larcombe, J. Lough, Temporal variation of light availability in coastal benthic habitats: Effects of clouds, turbidity, and tides. Limnology and Oceanography 49, 2201–2211 (2004).

52. S. DiPerna, M. Hoogenboom, S. Noonan, K. Fabricius, Effects of variability in daily light integrals on the photophysiology of the corals Pachyseris speciosa and Acropora millepora. PLoS ONE 13, e0203882 (2018).

53. P. G. Falkowski, J. A. Raven, Aquatic Photosynthesis (Princeton University Press, 2014).

54. T. López-Londoño, C. T. Galindo-Martínez, K. Gómez-Campo, L. A. González-Guerrero, S. Roitman, F. J. Pollock, V. Pizarro, M. López-Victoria, M. Medina, R. Iglesias-Prieto, Physiological and ecological consequences of the water optical properties degradation on reef corals. Coral Reefs 40, 1243–1256 (2021).

55. R. Hill, F. Bendall, Function of the Two Cytochrome Components in Chloroplasts: A Working Hypothesis. Nature 186, 136–137 (1960).

56. J. F. Hill, Govindjee, The controversy over the minimum quantum requirement for oxygen evolution. Photosynth Res 122, 97–112 (2014).

57. W. Skirving, S. Enríquez, J. D. Hedley, S. Dove, C. M. Eakin, R. A. B. Mason, J. L. D. L. Cour, G. Liu, O. Hoegh-Guldberg, A. E. Strong, P. J. Mumby, R. Iglesias-Prieto, Remote sensing of coral bleaching using temperature and light: Progress towards an operational algorithm. Remote Sensing 10, 18 (2018).

58. L. Muscatine, P. G. ; Falkowski, J. W. ; Porter, Z. Dubinsky, Fate of photosynthetic fixed carbon in light- and shade-adapted colonies of the symbiotic coral Stylophora pistillata. Proceedings of the Royal Society of London. Series B. Biological Sciences 222, 181–202 (1984).

59. L. Muscatine, Symbiosis of hydra and algae—III. Extracellular products of the algae. Comparative Biochemistry and Physiology 16, 77–92 (1965).

60. L. Muscatine, Glycerol excretion by symbiotic algae from corals and Tridacna and its control by the host. Science 156, 516–519 (1967).

61. M. S. Burriesci, T. K. Raab, J. R. Pringle, Evidence that glucose is the major transferred metabolite in dinoflagellate–cnidarian symbiosis. Journal of Experimental Biology 215, 3467–3477 (2012).

62. R. K. Trench, The Physiology and Biochemistry of Zooxanthellae Symbiotic with Marine Coelenterates. II. Liberation of Fixed 14C by Zooxanthellae in vitro. Proceedings of the Royal Society of London. Series B, Biological Sciences 177, 237–250 (1971).

63. R. K. Trench, The Physiology and Biochemistry of Zooxanthellae Symbiotic with Marine Coelenterates. I. The Assimilation of Photosynthetic Products of Zooxanthellae by Two Marine Coelenterates. Proceedings of the Royal Society of London. Series B, Biological Sciences 177, 225–235 (1971).

64. P. Tremblay, R. Grover, J. F. Maguer, M. Hoogenboom, C. Ferrier-Pagès, Carbon translocation from symbiont to host depends on irradiance and food availability in the tropical coral Stylophora pistillata. Coral Reefs 33, 1–13 (2014).

65. S. Sathyendranath, T. Platt, Ž. Kovač, J. Dingle, T. Jackson, R. J. W. Brewin, P. Franks, E. Marañón, G. Kulk, H. A. Bouman, Reconciling models of primary production and photoacclimation [Invited]. Applied Optics 59, C100 (2020).

66. R. D. Gates, P. J. Edmunds, The physiological mechanisms of acclimatization in tropical reef corals. American Zoologist 39, 30–43 (1999).

67. M. O. Hoogenboom, S. R. Connolly, K. R. N. Anthony, Effects of photoacclimation on the light niche of corals: A process-based approach. Marine Biology 156, 2493–2503 (2009).

68. B. Demmig-Adams, W. W. Adams, The role of xanthophyll cycle carotenoids in the protection of photosynthesis. Trends in Plant Science 1, 21–26 (1996).

69. B. E. Brown, I. Ambarsari, M. E. Warner, W. K. Fitt, R. P. Dunne, S. W. Gibb, D. G. Cummings, Diurnal changes in photochemical efficiency and xanthophyll concentrations in shallow water reef corals: Evidence for photoinhibition and photoprotection. Coral Reefs 18, 99–105 (1999).

70. P. J. Ralph, R. Gademann, A. W. D. Larkum, U. Schreiber, In situ underwater measurements of photosynthetic activity of coral zooxanthellae and other reef-dwelling dinoflagellate endosymbionts. Marine Ecology Progress Series 180, 139–147 (1999).

71. E. Erickson, S. Wakao, K. K. Niyogi, Light stress and photoprotection in Chlamydomonas reinhardtii. Plant Journal, 449–465 (2015).

72. R. J. Jones, O. Hoegh-Guldberg, Diurnal changes in the photochemical efficiency of the symbiotic dinoflagellates (Dinophyceae) of corals: photoprotection, photoinactivation and the relationship to coral bleaching. *Plant*, Cell & Environment 24, 89–99 (2001).

73. M. P. Lesser, M. Y. Gorbunov, Diurnal and bathymetric changes in chlorophyll fluorescence yields of reef corals measured in situ with a fast repetition rate fluorometer. 212, 69–77 (2001).

74. G. Winters, Y. Loya, R. Röttgers, S. Beer, Photoinhibition in shallow-water colonies of the coral Stylophora pistillata as measured in situ. Limnology and Oceanography 48, 1388–1393 (2003).

75. O. Levy, Z. Dubinsky, K. Schneider, Y. Achituv, D. Zakai, M. Y. Gorbunov, Diurnal hysteresis in coral photosynthesis. Marine Ecology Progress Series 268, 105–117 (2004).

76. A. Rodríguez-Román, X. Hernández-Pech, P. E. Thomé, S. Enríquez, R. Iglesias-Prieto, X. Herna, P. E. Thome, S. Enrı, R. Iglesias-Prieto, Photosynthesis and light utilization in the Caribbean coral Montastraea faveolata recovering from a bleaching event. Limnology and Oceanography 51, 2702–2710 (2006).

77. T. Scheufen, R. Iglesias-Prieto, S. Enríquez, Changes in the number of symbionts and Symbiodinium cell pigmentation modulate differentially coral light absorption and photosynthetic performance. Frontiers in Marine Science 4, 309 (2017).

78. T. Scheufen, W. E. Krämer, R. Iglesias-Prieto, S. Enríquez, Seasonal variation modulates coral sensibility to heat-stress and explains annual changes in coral productivity. Scientific Reports 7, 4937 (2017).

79. N. Kramer, R. Tamir, G. Eyal, Y. Loya, Coral Morphology Portrays the Spatial Distribution and Population Size-Structure Along a 5–100 m Depth Gradient. Front. Mar. Sci. 7 (2020).

80. J. P. Krall, G. E. Edwards, Relationship between photosystem II activity and CO2 fixation in leaves. Physiologia Plantarum 86, 180–187 (1992).

81. M. J. Fryer, J. R. Andrews, K. Oxborough, D. A. Blowers, N. R. Baker, Relationship between CO2 Assimilation, Photosynthetic Electron Transport, and Active O2 Metabolism in Leaves of Maize in the Field during Periods of Low Temperature1. Plant Physiology 116, 571–580 (1998).

82. O. Hoegh-Guldberg, R. Jones, Photoinhibition and photoprotection in symbiotic dinoflagellates from reef-building corals. Mar. Ecol. Prog. Ser. 183, 73–86 (1999).

83. M. Lesser, Depth-dependent photoacclimatization to solar ultraviolet radiation in the Caribbean coral Montastraea faveolata. Mar. Ecol. Prog. Ser. 192, 137–151 (2000).

84. J. A. Raven, The cost of photoinhibition. Physiologia Plantarum 142, 87–104 (2011).

85. X.-P. Yi, H.-S. Yao, D.-Y. Fan, X.-G. Zhu, P. Losciale, Y.-L. Zhang, W.-F. Zhang, W. S. Chow, The energy cost of repairing photoinactivated photosystem II: an experimental determination in cotton leaf discs. New Phytologist 235, 446–456 (2022).

86. S. Gunell, T. Lempiäinen, E. Rintamäki, E.-M. Aro, M. Tikkanen, Enhanced function of non-photoinhibited photosystem II complexes upon PSII photoinhibition. Biochimica et Biophysica Acta (BBA) - Bioenergetics 1864, 148978 (2023).

87. C. Langdon, On the causes of interspecific differences in the growth-irradiance relationship for phytoplankton. II. A general review. Journal of Plankton Research 10, 1291–1312 (1988).

88. N. Murata, S. Takahashi, Y. Nishiyama, S. I. Allakhverdiev, Photoinhibition of photosystem II under environmental stress. Biochimica et Biophysica Acta - Bioenergetics 1767, 414–421 (2007).

89. S. I. Allakhverdiev, Y. Nishiyama, S. Takahashi, S. Miyairi, I. Suzuki, N. Murata, Systematic Analysis of the Relation of Electron Transport and ATP Synthesis to the Photodamage and Repair of Photosystem II in Synechocystis. Plant Physiology 137, 263–273 (2005).

90. R. Iglesias-prieto, R. K. Trench, Acclimation and adaptation to irradiance in symbiotic dinoflagellates. 1.Responses of the photosynthetic unit to changes in photon flux density. Marine Ecology Progress Series 113, 163–176 (1994).

91. A. Trebst, Function of β-Carotene and Tocopherol in Photosystem II. Zeitschrift für Naturforschung C 58, 609–620 (2003).

92. A. Krieger-Liszkay, Singlet oxygen production in photosynthesis. J Exp Bot 56, 337–346 (2005).

93. M. E. Warner, W. K. Fitt, G. W. Schmidt, Damage to photosystem II in symbiotic dinoflagellates: a determinant of coral bleaching. Proceedings of the National Academy of Sciences of the United States of America 96, 8007–12 (1999).

94. S. Takahashi, S. Whitney, S. Itoh, T. Maruyama, M. Badger, Heat stress causes inhibition of the de novo synthesis of antenna proteins and photobleaching in cultured Symbiodinium. Proceedings of the National Academy of Sciences of the United States of America 105, 4203–8 (2008).

95. S. Takahashi, M. Yoshioka-Nishimura, D. Nanba, M. R. Badger, Thermal Acclimation of the Symbiotic Alga Symbiodinium spp. Alleviates Photobleaching under Heat Stress. PLANT PHYSIOLOGY 161, 477–485 (2013).

96. R. Hill, C. M. Brown, K. DeZeeuw, D. a Campbell, P. J. Ralph, Increased rate of D1 repair in coral symbionts during bleaching is insufficient to counter accelerated photo-inactivation. Limnology and Oceanography 56, 139–146 (2011).

97. S. Takahashi, S. E. Milward, W. Yamori, J. R. Evans, W. Hillier, M. R. Badger, The solar action spectrum of photosystem II damage. Plant Physiology 153, 988–993 (2010).

98. J. M. Pandolfi, C. E. Lovelock, A. F. Budd, Character Release Following Extinction in a Caribbean Reef Coral Species Complex. Evolution 56, 479–501 (2002).

99. C. Prada, M. Gómez-Corrales, A. González, B. Kamel, M. Medina, N. Knowlton, D. Levitan, Speciation across depth gradients in reef corals. In Review [Preprint] (2024). 10.21203/rs.3.rs-4202515/v1.

100. H. E. Aichelman, B. E. Benson, K. Gomez-Campo, M. I. Martinez-Rugerio, J. E. Fifer, L. Tsang, A. M. Hughes, C. B. Bove, O. C. Nieves, A. M. Pereslete, D. Stanizzi, N. G. Kriefall, J. H. Baumann, J. P. Rippe, P. Gondola, K. D. Castillo, S. W. Davies, Cryptic coral diversity is associated with symbioses, physiology, and response to thermal challenge. Science Advances 11, eadr5237 (2025).

101. D. Iluz, I. Alexandrovich, Z. Dubinsky, The enhancement of photosynthesis by fluctuating light. Artifical Photosynthesis (2012).

102. J. E. Faust, V. Holcombe, N. C. Rajapakse, D. R. Layne, The effect of daily light integral on bedding plant growth and flowering. HortScience 40, 645–649 (2005).

103. S. W. Jeffrey, G. F. Humphrey, New spectrophotometric equations for determining chlorophylls a, b, c1 and c2 in higher plants, algae and natural phytoplankton. Biochemie und Physiologie der Pflanzen 167, 191–194 (1975).

104. R. M. Vásquez-Elizondo, L. Legaria-Moreno, M. Á. Pérez-Castro, W. E. Krämer, T. Scheufen, R. Iglesias-Prieto, S. Enríquez, Absorptance determinations on multicellular tissues. Photosynthesis Research 132, 311–324 (2017).

105. S. Enríquez, K. Sand-Jensen, Variation in Light Absorption Properties of *Mentha aquatica* L. as a Function of Leaf Form: Implications for Plant Growth. International Journal of Plant Sciences 164, 125–136 (2003).

106. N. M. Cayabyab, S. Enríquez, Leaf photoacclimatory responses of the tropical seagrass *Thalassia testudinum* under mesocosm conditions: a mechanistic scaling-up study. New Phytologist 176, 108–123 (2007).

107. L. A. González-Guerrero, R. M. Vásquez-Elizondo, T. López-Londoño, G. Hernán, R. Iglesias-Prieto, S. Enríquez, Validation of parameters and protocols derived from chlorophyll a fluorescence commonly utilised in marine ecophysiological studies. Functional Plant Biol., doi: 10.1071/FP21101 (2021).

108. W. C. Dennison, R. S. Alberte, Photosynthetic Responses of Zostera marina L. (Eelgrass) to in situ Manipulations of Light Intensity. Oecologia 55, 137–144 (1982).

109. M. K. DeSalvo, S. Sunagawa, P. L. Fisher, C. R. Voolstra, R. Iglesias-Prieto, M. Medina, Coral host transcriptomic states are correlated with Symbiodinium genotypes. Molecular Ecology 19, 1174–1186 (2010).

110. D. W. Kemp, D. J. Thornhill, R. D. Rotjan, R. Iglesias-Prieto, W. K. Fitt, G. W. Schmidt, Spatially distinct and regionally endemic Symbiodinium assemblages in the threatened Caribbean reef-building coral Orbicella faveolata. Coral Reefs 34, 535–547 (2015).

111. A. M. Lewis, A. N. Chan, T. C. LaJeunesse, New Species of Closely Related Endosymbiotic Dinoflagellates in the Greater Caribbean have Niches Corresponding to Host Coral Phylogeny. Journal of Eukaryotic Microbiology 66, 469–482 (2019).

112. T. C. LaJeunesse, W. K. W. Loh, R. Van Woesik, O. Hoegh-Guldberg, G. W. Schmidt, W. K. Fitt, Low symbiont diversity in southern Great Barrier Reef corals, relative to those of the Caribbean. Limnology and Oceanography 48, 2046–2054 (2003).

113. T. C. Lajeunesse, R. K. Trench, Biogeography of two species of Symbiodinium (Freudenthal) inhabiting the intertidal sea anemone Anthopleura elegantissima (Brandt). 10.2307/1542872 199, 126–134 (2016).

114. R. Zardoya, E. Costas, V. López-Rodas, A. Garrido-Pertierra, J. M. Bautista, Revised dinoflagellate phylogeny inferred from molecular analysis of large-subunit ribosomal RNA gene sequences. Journal of Molecular Evolution 41, 637–645 (1995).

115. H. Li, Aligning sequence reads, clone sequences and assembly contigs with BWA-MEM. arXiv arXiv:1303.3997 [Preprint] (2013). 10.48550/arXiv.1303.3997.

116. V. Avila-Magaña, B. Kamel, M. DeSalvo, K. Gómez-Campo, S. Enríquez, H. Kitano, R. V. Rohlfs, R. Iglesias-Prieto, M. Medina, Elucidating gene expression adaptation of phylogenetically divergent coral holobionts under heat stress. Nat Commun 12, 5731 (2021).

117. E. F. Camp, T. Kahlke, B. Signal, C. A. Oakley, A. Lutz, S. K. Davy, D. J. Suggett, W. P. Leggat, Proteome metabolome and transcriptome data for three Symbiodiniaceae under ambient and heat stress conditions. Sci Data 9, 153 (2022).

118. R. A. Levin, V. H. Beltran, R. Hill, S. Kjelleberg, D. McDougald, P. D. Steinberg, M. J. H. van Oppen, Sex, Scavengers, and Chaperones: Transcriptome Secrets of Divergent Symbiodinium Thermal Tolerances. Mol Biol Evol 33, 2201–2215 (2016).

119. C. M. Brown, D. A. Campbell, J. E. Lawrence, Resource dynamics during infection of Micromonas pusilla by virus MpV-Sp1. Environmental Microbiology 9, 2720–2727 (2007).

120. C. Six, Z. V. Finkel, A. J. Irwin, D. A. Campbell, Light Variability Illuminates Niche-Partitioning among Marine Picocyanobacteria. PLoS ONE 2, e1341 (2007).

121. P. Ewels, M. Magnusson, S. Lundin, M. Käller, MultiQC: summarize analysis results for multiple tools and samples in a single report. Bioinformatics 32, 3047–3048 (2016).

122. A. M. Bolger, M. Lohse, B. Usadel, Trimmomatic: a flexible trimmer for Illumina sequence data. Bioinformatics 30, 2114–2120 (2014).

123. N. L. Bray, H. Pimentel, P. Melsted, L. Pachter, Near-optimal probabilistic RNA-seq quantification. Nat Biotechnol 34, 525–527 (2016).

124. L. Fu, B. Niu, Z. Zhu, S. Wu, W. Li, CD-HIT: accelerated for clustering the next-generation sequencing data. Bioinformatics 28, 3150–3152 (2012).

125. C. Camacho, G. Coulouris, V. Avagyan, N. Ma, J. Papadopoulos, K. Bealer, T. L. Madden, BLAST+: architecture and applications. BMC Bioinformatics 10, 421 (2009).

126. J. Kirk, Light and Photosynthesis (Cambridge University Press, New York, NY, 2011; https://www.cambridge.org/core/books/light-and-photosynthesis-in-aquatic-ecosystems/C19B28AE07B1CDEBDA5593194DE4E304).

127. A. D. Jassby, T. Platt, Mathematical formulation of the relationship between photosynthesis and light for phytoplankton. Limnology and Oceanography 21, 540–547 (1976).

