## Supplementary Materials for "Maintenance cost of photosynthesis sets key ecological constraints on zooxanthellate corals"

Gomez-Campo *et al.*

**This PDF file includes:**

Figs. S1 to S7
Table S1

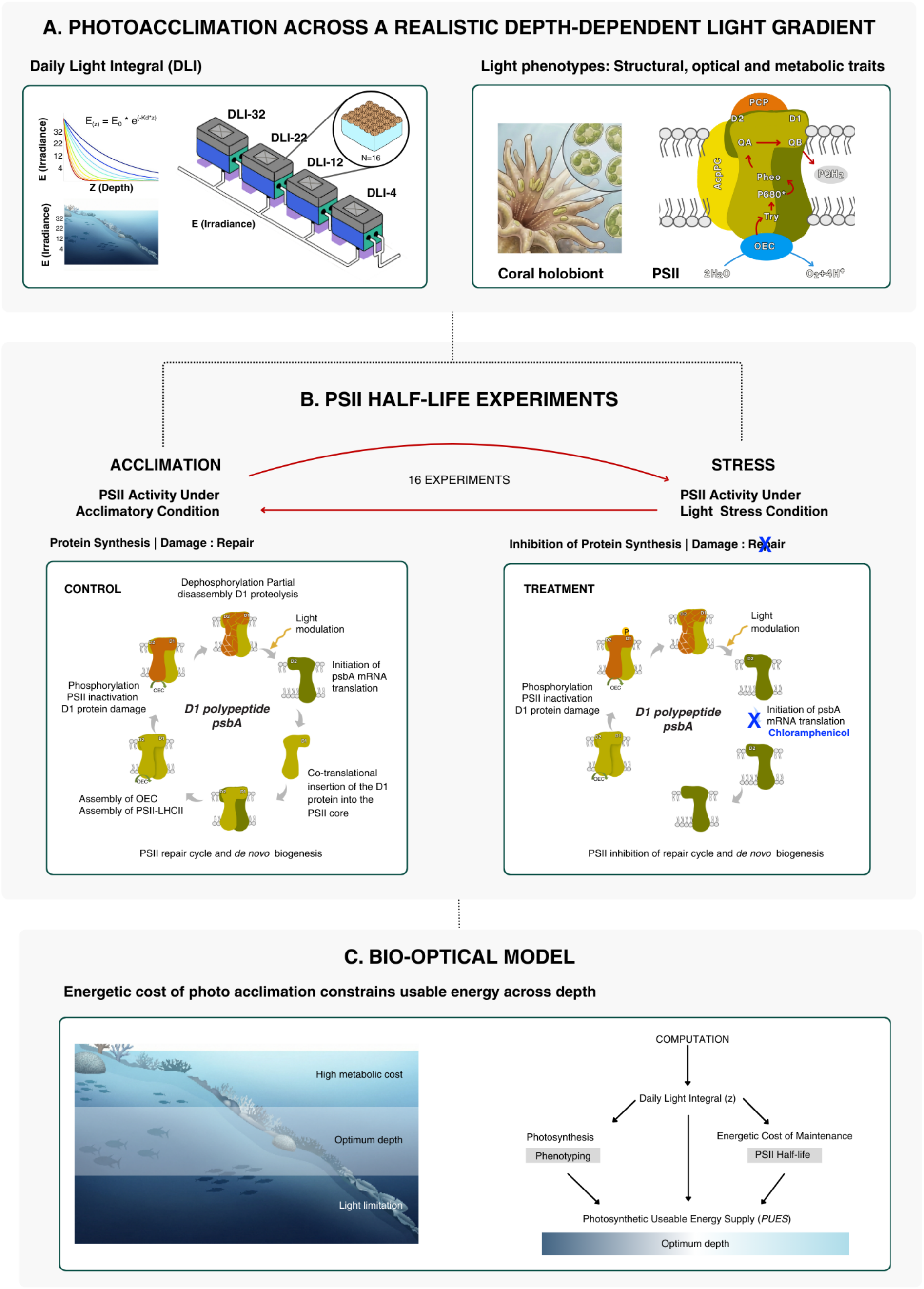


**Fig. S1.** Experimental approach linking light environment, PSII repair dynamics, and energetic modeling. (**A**) Corals were acclimated to four controlled daily light integrals (DLI-4, -12, -22, -32 mol photons $m^{-2}$ $d^{-1}$) based on a realistic depth gradient (based on $K_d$) and generated by programmable LEDs simulating diurnal cycles to characterize light-dependent phenotypes and PSII activity. (**B**) PSII repair activity were quantified after acclimation and in light stress (switch to different DLIs) by inhibiting D1 synthesis and estimating PSII half-life ($t_1/_2$) from hourly photochemical efficiency measurements. (**C**) A depth-explicit bio-optical model integrates light attenuation, photosynthetic performance, and PSII turnover to estimate Photosynthetic Usable Energy Supply (PUES) and identify the energetic optimum.

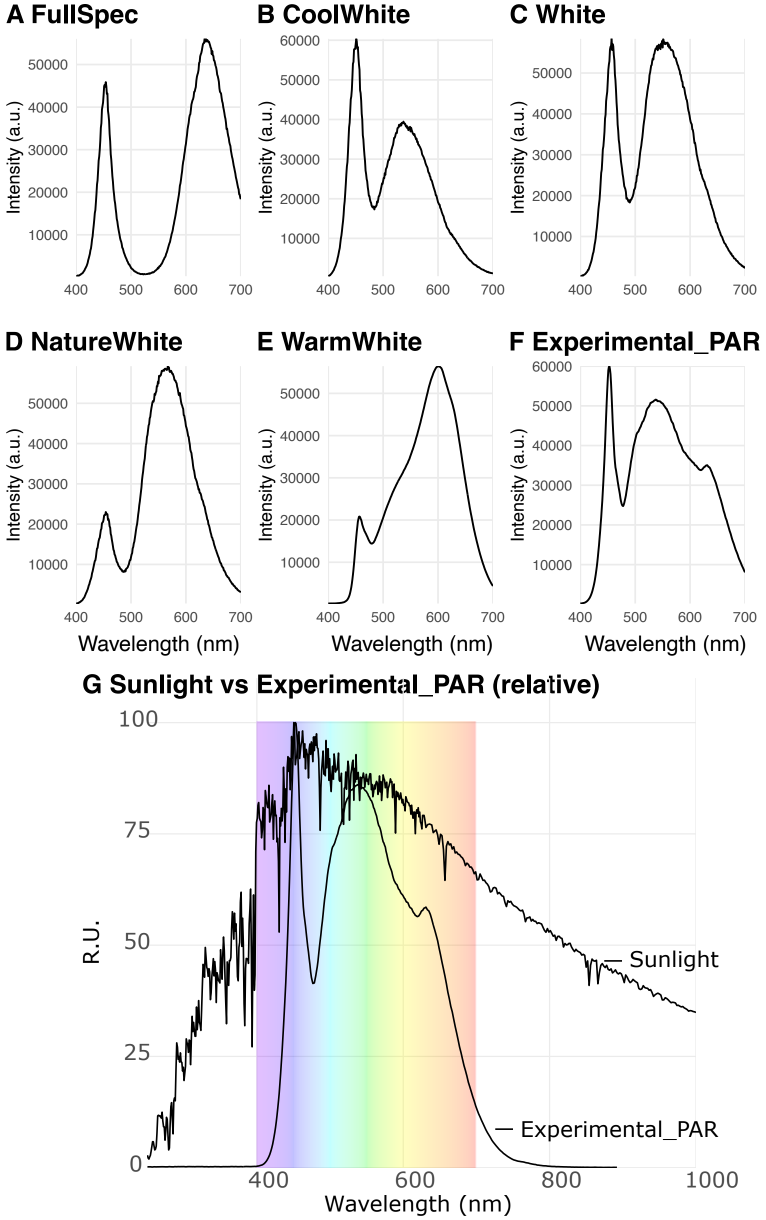


**Fig. S2. Spectral characterization of the experimental light field. (A–F)** Emission spectra of commercial LEDs tested and (**G**) spectrum (experimental_PAR) delivered to corals under experimental conditions, measured with an Ocean Optics USB2000+ spectrometer. The colored background denotes the photosynthetically active radiation (PAR; 400–700 nm), illustrating that the assembled light field approximates a broad, sun-like spectral distribution within the PAR region. Spectra are normalized to their respective maxima for visualization.

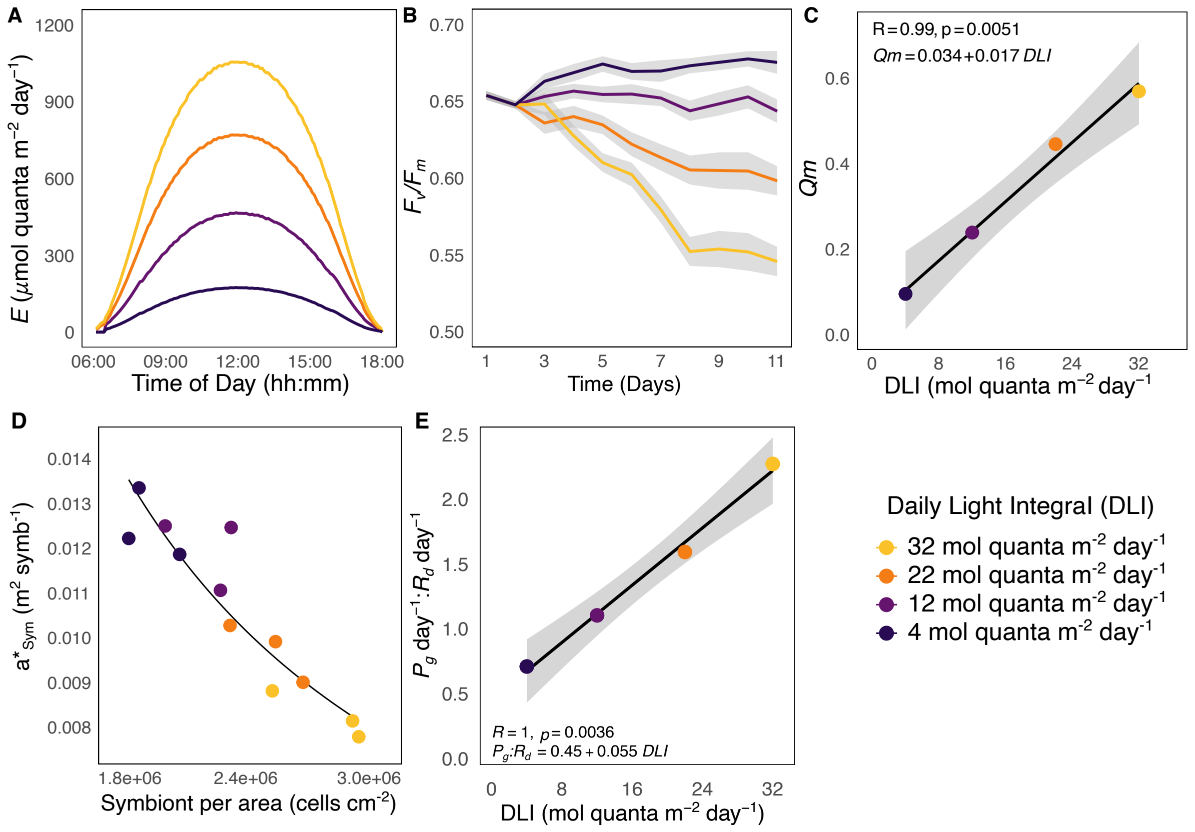


**Figure S3**. **Phenotyping after acclimation to light conditions.** (**A**) Calibrated light conditions, measured, monitored with the quantum sensor LI-1400 (LI-COR, USA). (**B**) Acclimatory kinetics of maximum photochemical efficiency of PSII ($F_v/F_m$). **(C)** PSII excitation pressure ($Q_m$) after 4$^+$ weeks of acclimation, illustrating a linear trend with light exposure and increased symbiont contribution to host metabolism under higher light. Symbols represent mean ± SE (n = 16). **(D)** Symbiont-specific absorption ($a^*_{Sym}$) showing optimization of symbiont cell absorption efficiency as light decreases. (**E)** Daily integrated photosynthesis-to-respiration ratios, an index of autotrophic capacity. Symbols represent mean ± SE (n = 8). Shaded areas show bootstrapped 95% confidence intervals.

A

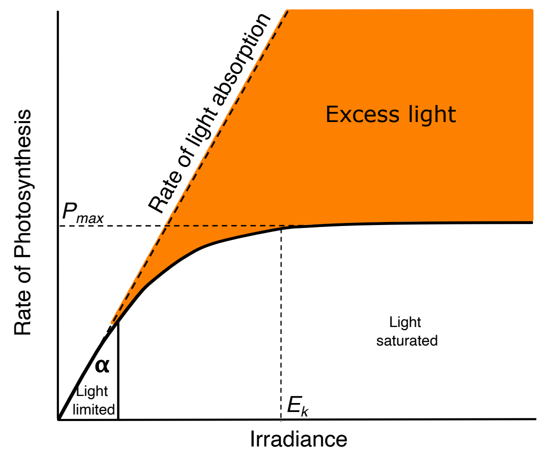


B

| Descriptor | Definition | Function | Type | p-value |
| --- | --- | --- | --- | --- |
| **Photochemical efficiency of PSII** | | | | |
| *Fv/Fm* | Maximum photochemical efficiency. | $Fv/_{Fm} = 0.7 - 0.0045\ DLI$ | Linear regression | p = 0.005 |
| *Qm* | Maximum excitation pressure over PSII. | $Qm = 0.034 + 0.017\ DLI$ | Linear regression | p = 0.005 |
| **PSII reaction center - D1 molecular activity** | | | | |
| *PSII1/2* | PSII lifetime | $PSII_{(1/2,h)} = 36.38\ DLI^{-0.72}$ | Power functions | p = 0.013 |
| *D1* protein-1 | *D1* protein abundance per unit of total protein | $D1 = 0.804\ DLI^{-0.734}$ | Power function | p = 0.028 |
| *D1* Chla-1 | *D1* protein abundance per unit Chla | $D1 = 41.980\ DLI^{-0.511}$ | Power function | p = 0.035 |
| *D1* ~ PSII1/2 | D1 protein-1 ~ PSII lifetime | $D1 = -0.002 + 0.022 PSII_{1/2}$ | Linear regression | p = 0.001 |
| **Optical properties of the host tissue** | | | | |
| Chla | Chla abundance | $Chla = 212.54\ DLI^{-0.45}$ | Power function | p < 0.001 |
| *a*Chla* | Light absorption efficiency per unit of Chla | $a*_{Chla} = 1.537\ Chla^{-0.883}$ | Power function | p < 0.001 |
| **Metabolic rates** | | | | |
| *α* | Initial slope of the P–E curve. | $\alpha = 0.0362 - 0.0005\ DLI$ | Linear regression | p = 0.006 |
| *Ec* | Compensation irradiance | $E_c = 58.1 + 3.92\ DLI$ | Linear regression | p = 0.029 |
| *Ek* | Saturation irradiance | $Ek = 300 + 9.5\ DLI$ | Linear regression | p = 0.008 |
| Rd | Dark respiration rate of the coral holobiont | $R_d = 2.8 + 0.025\ DLI$ | Linear regression | p = 0.006 |

C

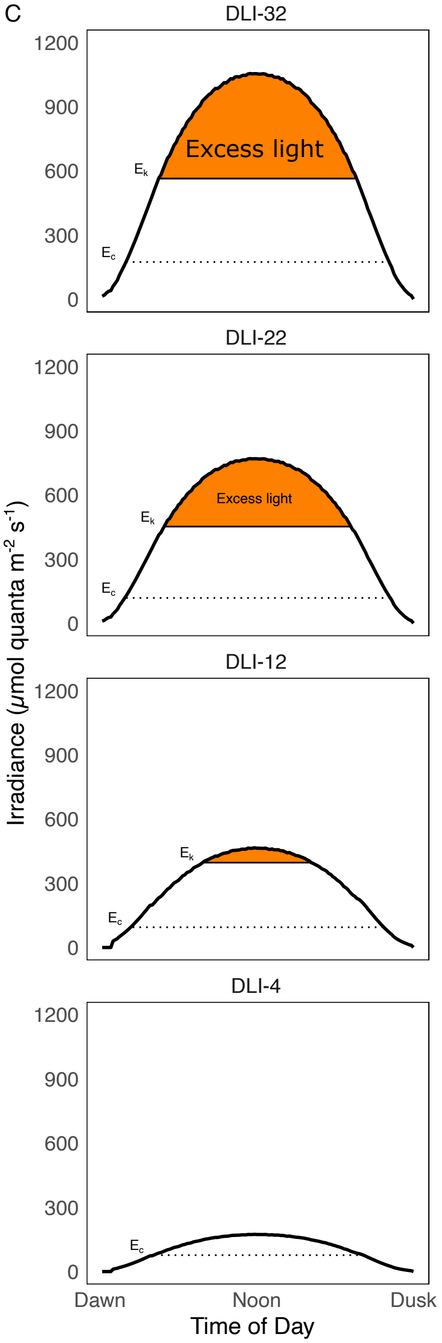


**Fig. S4. Function-valued phenotyping of across daily light integrals. (A)** Typical photosynthesis-irradiance (P-E) curve illustrating the transition from light limitation to light saturation and the accumulation of excess absorbed light at high irradiance. (**B**) Summary of functional traits, fitted equations, and their relationship with daily light integral (DLI). (**C**) Diurnal trajectories of photosynthetic descriptors. Orange shading indicates periods in which absorbed irradiance no longer results in more carbon fixation ("excess light"), dotted horizontal line are periods at compensation irradiance ($E_c$) and continuous line periods at saturation irradiance ($E_k$).

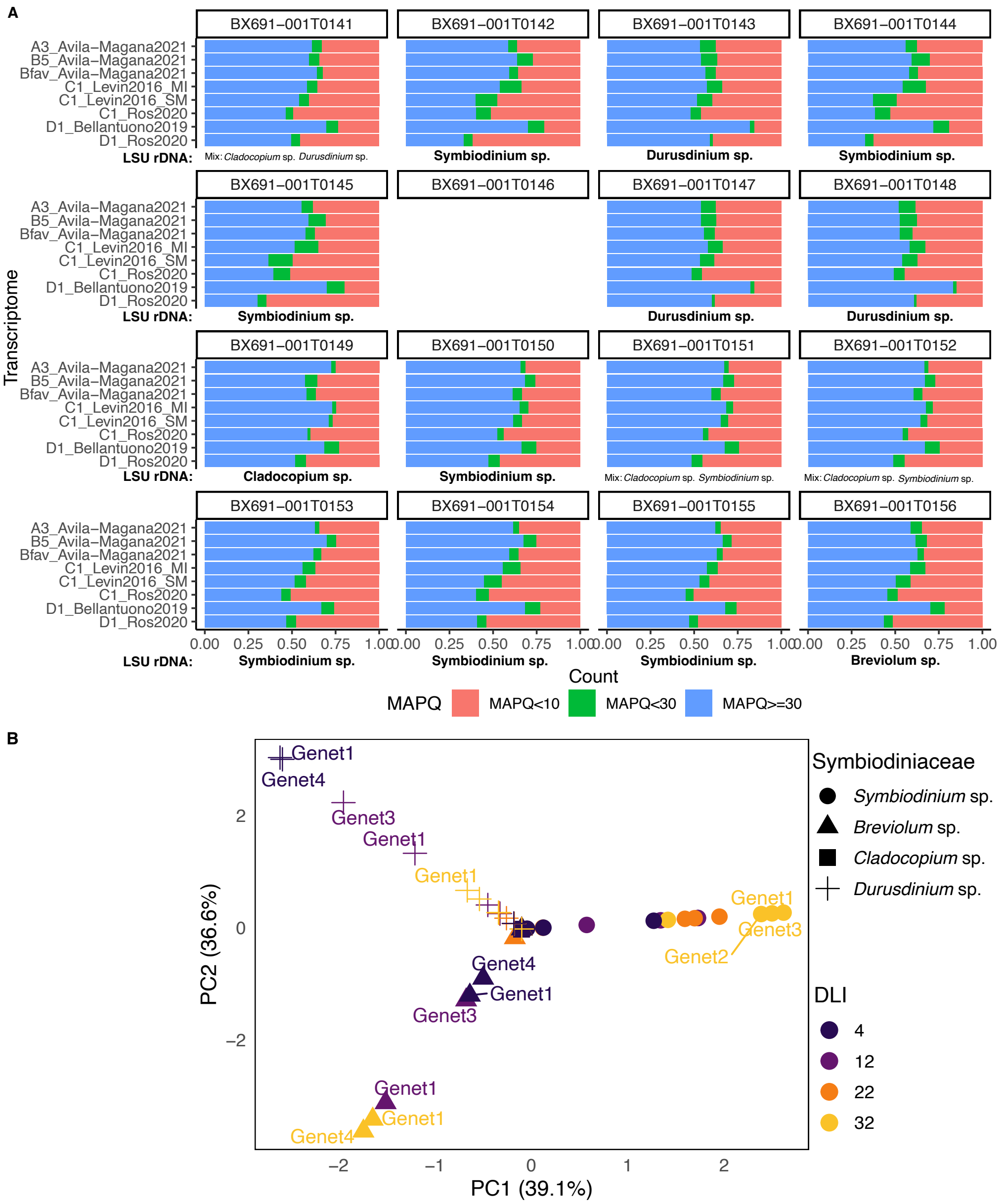


**Fig. S5. Symbiont typing of experimental colonies using qPCR and transcriptomic signatures.** (**A**) Symbiont community profiles for each colony, assigned by the large ribosomal large subunit (*LSU* rDNA) and cross-validated by mapping RNA-seq reads to published Symbiodiniaceae reference transcriptomes (see methods). Samples used in this section (n = 4) are arranged by light condition from minimum (top) to maximum (bottom) irradiance. Colony with missing data is indicated by a blank panel. (**B**) Principal component analysis (PCA) of *psbA* transcript expression across all colonies. Each point represents a colony, labeled with its genet ID and annotated by symbiont genus (symbol) and experimental light condition (color). PC1 and PC2 explained 75.7% of the total variance. Clustering by Symbiodiniaceae genus was visible (along PC1), showing PC1 likely captures symbiont genus-specific expression of *psbA* gene, and suggesting core differences in expression profiles among genera. DLI regulation was not clearly reflected. Colony genet identity did not dominate structure, implying host background does not explain most variation in *psbA* expression.

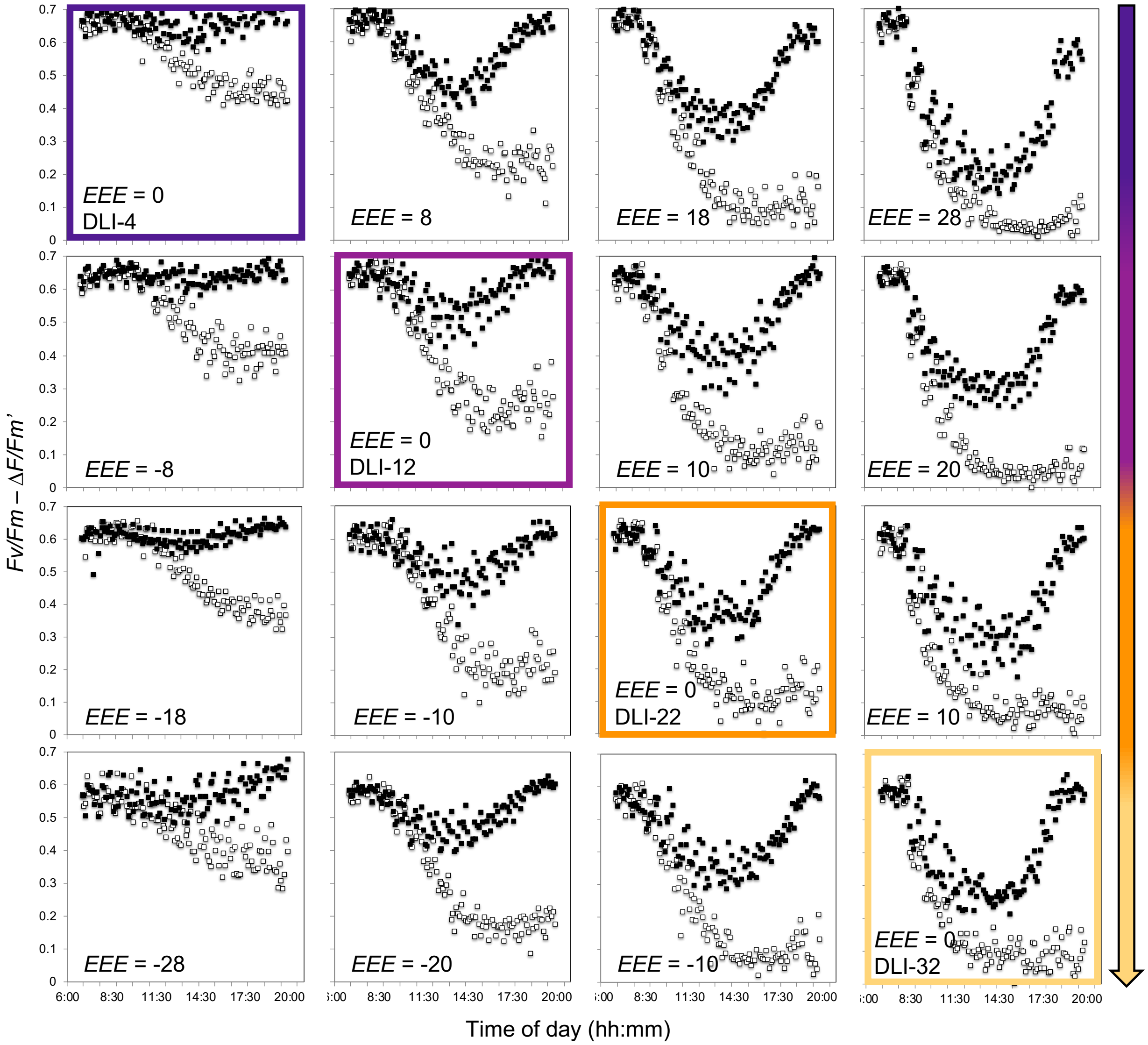


**Fig. S6. Diurnal PSII activity under light stress conditions.** Comparison of the severity of light stress in *Orbicella faveolata*. (A) After 4+ weeks of acclimation to each DLI (colored boxes), corals were switched to alternative light exposures for one diurnal cycle. Excess excitation energy (EEE, mol quanta $m^2$ $d^{-1}$) was estimated as the difference between DLI-destination and DLI-acclimation. Each row represents the change in light condition for each acclimatory state, *i.e.*, first row is DLI-4 (EEE = 0 colored panel, corals acclimated to DLI-4) switched to DLI-12 (EEE = 8), DLI-22 (EEE = 18) and DLI-32 (EEE = 28). Monitoring of photochemical efficiency showed photoinactivation and repair cycles in PSII in the presence (empty squares) and absence (filled squares) of protein synthesis inhibitor chloramphenicol (CAP).

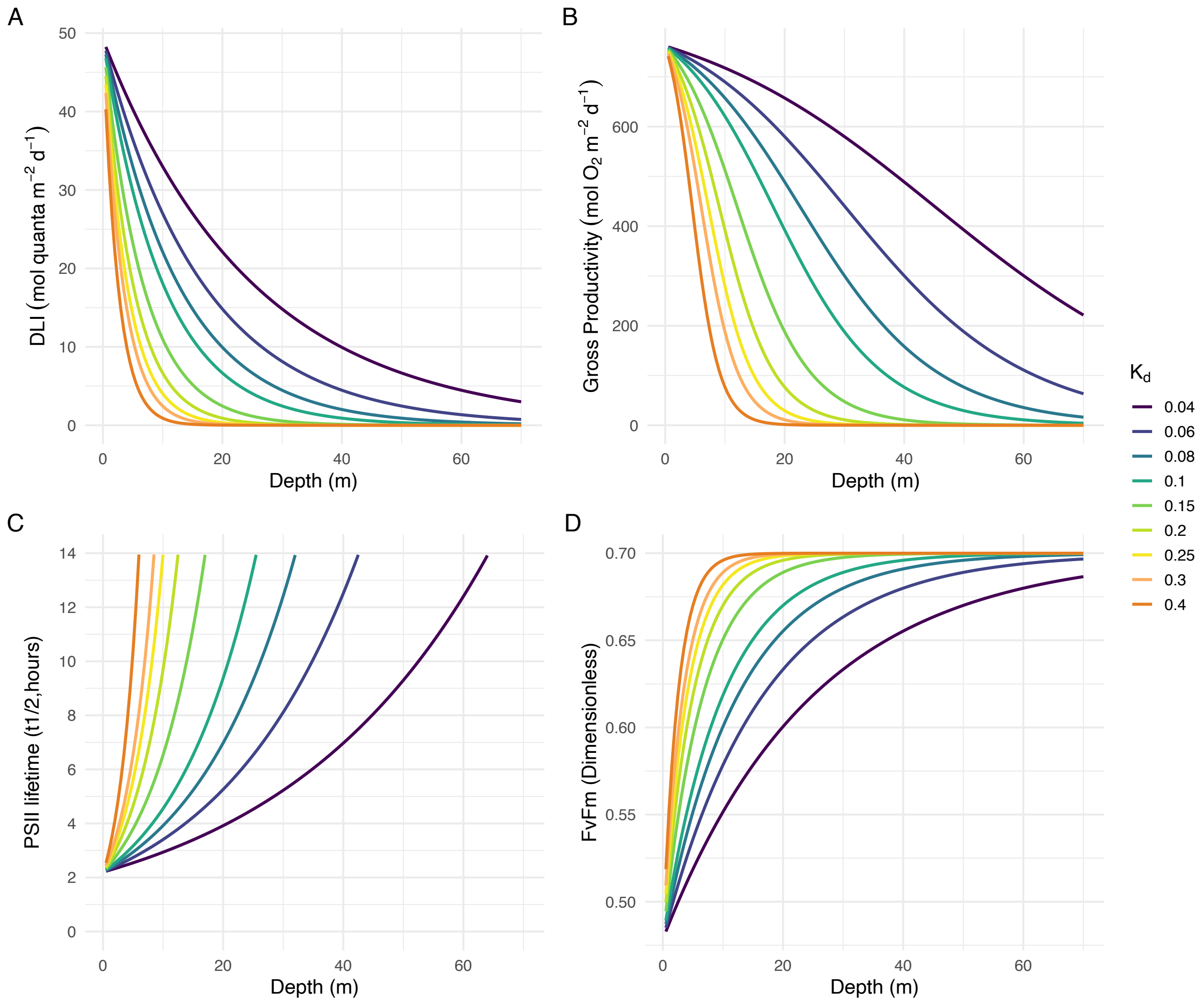


**Fig. S7. Depth-dependent bio-optical modeling of irradiance, photosynthetic performance, and PSII repair dynamics.** (**A**) Daily light integral (DLI) as a function of depth for a range of diffuse attenuation coefficients ($K_d$ = 0.04-0.40 $m^{-1}$), computed from the exponential decay of downwelling irradiance. Curves represent contrasting optical water types from very clear (low Kd) to highly turbid (high Kd) conditions. (**B**) Modeled daily gross photosynthetic output (GPg(z)) across depths, derived by integrating photosynthesis-irradiance (P-E) relationships over the diel light cycle using the DLI profiles in panel A. (**C**) Estimated PSII reaction-center lifetime ($t_{1/2}$) as a function of depth, based on the empirically derived relationship between DLI and D1 turnover. Increasing $t_{1/2}$ at depth reflects reduced requirements for PSII repair under low-irradiance conditions. (**D**) Modeled maximum photochemical efficiency ($F_v/F_m$) as a function of depth, calculated from the fitted DLI-$F_v/F_m$ function. Main descriptors used to parameterize the bio-optical model used to compute the Photosynthetic Usable Energy Supply (*PUES*) in Fig. 6.

**Table S1. Table of results.** Phenotyping of coral holobionts.

| Units | Parameters and notations | Units | DLI-4 | | DLI-12 | | DLI-22 | | DLI-32 | |
|---|---|---|---|---|---|---|---|---|---|---|
| | | | Mean | SE | Mean | SE | Mean | SE | Mean | SE |
| PSII | Maximum photochemical efficiency ($F_v/F_m$) | Dimensionless | 0.6753 | 0.0074 | 0.6435 | 0.0076 | 0.5982 | 0.0095 | 0.5457 | 0.0095 |
| | Effective photochemical efficiency ($\Delta F/Fm'$) | Dimensionless | 0.6100 | 0.0058 | 0.4911 | 0.0150 | 0.3427 | 0.0194 | 0.2373 | 0.0135 |
| | Maximum excitation pressure over PSII ($Q_m$) | Dimensionless | 0.0825 | 0.0062 | 0.2545 | 0.0211 | 0.4684 | 0.0211 | 0.5860 | 0.0233 |
| | PSII half-life ($PSII_{1/2}$) | Hours | 13.414 | 0.015 | 5.690 | 0.203 | 4.521 | 0.255 | 2.698 | 1.215 |
| | PSII D1 protein abundance per protein | pmol D1 µg total protein$^{-1}$ | 0.2929 | 0.0311 | 0.1142 | 0.0233 | 0.1040 | 0.0173 | 0.0576 | 0.0050 |
| | PSII D1 protein abundance per Chl*a* | pmol D1 µg Chla$^{-1}$ | 20.341 | 3.952 | 12.759 | 0.758 | 9.358 | 2.510 | 5.561 | 0.391 |
| Structural and optical traits | Chl*a* density per area | mg Chl*a* m$^{-2}$ | 111.534 | 7.237 | 76.776 | 7.515 | 49.086 | 5.654 | 41.153 | 3.966 |
| | Zoox. per area of host tissue | Cells cm$^2$ | 1,973,601 | 73,384 | 2,239,944 | 96,497 | 2,541,594 | 100,458 | 2,817,193 | 131,434 |
| | Chl*a* per zoox. cell (*Ci*) | Pg Chl*a* sym$^{-1}$ | 5.6845 | 0.5183 | 3.4241 | 0.2755 | 1.9379 | 0.2310 | 1.4691 | 0.1601 |
| | Absorptance PAR (*APAR*) | Dimensionless | 0.8932 | 0.0062 | 0.8844 | 0.0074 | 0.8663 | 0.0094 | 0.8366 | 0.0123 |
| | Absorptance 675 (A675) | Dimensionless | 0.9249 | 0.0036 | 0.9255 | 0.0048 | 0.9158 | 0.0058 | 0.8992 | 0.0086 |
| | Absorbance PAR (*DePAR*) | Dimensionless | 1.0083 | 0.0238 | 0.9850 | 0.0284 | 0.9237 | 0.0277 | 0.8482 | 0.0337 |
| | Absorbance 675 (*De*675) | Dimensionless | 1.1334 | 0.0206 | 1.1268 | 0.0271 | 1.0809 | 0.0272 | 1.0070 | 0.0358 |
| | Chla specific absorption coefficient (a*Chla) | $m^2$ mg Chl*a*$^{-1}$ | 0.0222 | 0.0015 | 0.0357 | 0.0040 | 0.0513 | 0.0047 | 0.0573 | 0.0054 |
| | Symbiont specific absorption coefficient ($a^*_{Sym}$) | $m^2$ sym$^{-1}$ | 0.0125 | 0.0004 | 0.0120 | 0.0005 | 0.0097 | 0.0004 | 0.0082 | 0.0003 |
| Metabolic rates | Photosynthetic efficiency ($\alpha$) | mol $O_2$ mol incident quanta$^{-1}$ | 0.0336 | 0.0018 | 0.0308 | 0.0007 | 0.0257 | 0.0013 | 0.0196 | 0.0018 |
| | Quantum yield of photosynthesis ($\Phi_{O2}$) | mol $O_2$ mol absorbed quanta$^{-1}$ | 0.1059 | 0.0022 | 0.0906 | 0.0039 | 0.0803 | 0.0044 | 0.0589 | 0.0057 |
| | Minimum Quantum Requirement ($1/\Phi_{O2}$) | photons absorbed per $O_2$ evolved | 9.425 | 0.197 | 11.220 | 0.481 | 12.690 | 0.648 | 17.338 | 1.893 |
| | Maximum photosynthetic rate (*Pmax*) | µmol $O_2$ $m^{-2}$ $s^{-1}$ | 11.906 | 0.569 | 13.018 | 0.936 | 12.351 | 0.538 | 11.463 | 0.470 |
| | Saturation irradiance ($E_k$) | µmol quanta $m^{-2}$ $s^{-1}$ | 341.618 | 12.617 | 420.870 | 24.549 | 486.988 | 28.039 | 616.823 | 53.883 |

| | | | | | | | | | |
|---|---|---|---|---|---|---|---|---|---|
| Compensation irradiance ($E_c$) | µmol quanta $m^{-2}$ $s^{-1}$ | 82.226 | 5.809 | 100.726 | 10.094 | 129.535 | 7.566 | 194.200 | 18.455 |
| Dark respiration (*Rd*) | µmol $O_2$ $m^{-2}$ $s^{-1}$ | -2.3314 | 0.1639 | -2.4782 | 0.1148 | -2.4177 | 0.1142 | -2.9655 | 0.1681 |
| Light-enhanced respiration ($R_l$) | µmol $O_2$ $m^{-2}$ $s^{-1}$ | -3.3916 | 0.3187 | -3.7038 | 0.5434 | -4.1160 | 0.1448 | -4.2208 | 0.2115 |
| Dark respiration ($R_D$; avg Rd Rl) | µmol $O_2$ $m^{-2}$ $s^{-1}$ | 2.8615 | 0.2187 | 3.0910 | 0.2863 | 3.2669 | 0.0879 | 3.5931 | 0.1704 |
| Photosynthetic rate per zoox. cell (*Psym*) | µmol $O_2$ $m^{-2}$ $s^{-1}$ | 2.0209 | 0.2651 | 2.0082 | 0.1516 | 1.6124 | 0.0946 | 1.5214 | 0.2280 |
